# Physiological folate levels constrain nucleotide synthesis and increase dependence on nucleotide salvage

**DOI:** 10.64898/2026.09.13.750663

**Authors:** Ryan Elbashir, Keene L. Abbott, Diya L. Ramesh, Ahmed Ali, Anna Shevzov-Zebrun, Anna M. Barbeau, Michelle Wu, Abigail P. Ward, Yetiş Gültekin, Brian T. Do, Sharanya Sivanand, Azucena Ramos, Tenzin Kunchok, Millenia Waite, Edrees H. Rashan, Muhammad Bin Munim, Michael T. Hemann, Matthew G. Vander Heiden

## Abstract

Proliferating cells must acquire nucleotides to support DNA replication, yet how cells meet these nucleotide demands for proliferation under physiological conditions remains understudied. Here, we investigated how physiological nutrient availability shapes nucleotide acquisition strategies in a mouse model of B-cell acute lymphoblastic leukemia (B-ALL). To assess how environmental nutrients impact nucleotide metabolism, we formulated a mouse plasma-like medium (MPM) that reproduces the circulating metabolite composition of plasma from mice with B-ALL and assessed how this influenced nucleotide metabolism relative to standard culture conditions, where nucleotide acquisition has historically been studied. We find that leukemia cells cultured in MPM acquire nucleotides through salvage pathways, and that select nucleotide salvage pathways are required for proliferation under physiological conditions. Of note, this dependency on nucleotide salvage in plasma-like conditions was not caused by precursor metabolite limitation for *de novo* synthesis. Instead, we found that physiological folate levels are insufficient to support deoxynucleotide triphosphate (dNTP) synthesis for genome replication, leading to DNA replication stress and impaired proliferation when nucleotide salvage is disrupted. Consistently, dietary folate restriction exacerbates the impaired leukemia progression phenotype of nucleotide salvage-deficient B-ALL cells. Together, these findings demonstrate that access to folates is an endogenous limitation for nucleotide synthesis in plasma-like nutrient conditions, increasing the relevance of nucleotide salvage pathways for leukemia progression. More broadly, this work highlights how micronutrient abundance can influence metabolic dependencies and reveals that folate levels shape nucleotide metabolism under physiological conditions.

## Introduction

Proliferating cells reprogram metabolism to meet the biosynthetic demands of cell growth and division^1^. Among the macromolecules required for cell proliferation, the acquisition of nucleotides is often limiting. As a consequence, failure to maintain the appropriate levels and balance of dNTPs can impair DNA replication and cell cycle progression^2–4^. In proliferating mammalian cells, dNTP levels are tightly regulated and increase during S phase to ensure a sufficient supply for genome duplication^5^, and disruption of dNTP availability or in the balance between individual dNTP species leads to DNA replication stress, which can progress to DNA damage and proliferative arrest^3,5^. Thus, understanding how cells source the nucleotides needed for dNTP production and DNA replication is essential to understand proliferative cell metabolism.

Eukaryotic cells acquire nucleotides through three principal routes: *de novo* synthesis, salvage of extracellular nucleobases and nucleosides, and recycling from the degradation of intracellular nucleic acid-containing material such as ribosomes^4^. *De novo* nucleotide synthesis requires access to multiple precursor metabolites, including carbohydrates to generate ribose, and the amino acids serine, glycine, glutamine, and aspartate, as well as vitamins in the form of folates and electron carriers like nicotinamide adenine dinucleotide (NAD^+^) and nicotinamide adenine dinucleotide phosphate (NADPH) to maintain redox balance^4,6^. Prior work has demonstrated that limiting redox balance, or access to substrates such as serine or aspartate, can limit *de novo* nucleotide synthesis and restrict proliferation, suggesting nucleotide production is a metabolic bottleneck for cells in some environmental conditions^6–8^.

The availability of extracellular nucleotide precursors adds another layer of complexity to how cells meet their nucleotide demands. When available, nucleotide precursors can be salvaged from the extracellular environment. Nucleotides, nucleosides, and nucleobases are present at varying levels across tissues and disease states^9–11^, and nucleotide-related metabolites account for a substantial proportion of the variance in metabolite levels between anatomical sites^9^. Yet, most mechanistic studies of nucleotide metabolism *in vitro* have been conducted in standard cell culture media, which lack nucleotides and products of nucleotide metabolism^12^. Numerous studies have shown that nucleotide salvage pathways are engaged when precursors are available or when synthesis is inhibited^13–15^, and recent studies in mice have reported that salvage pathways can contribute to nucleotide levels in both normal and malignant tissues^10^. While this work confirms salvage pathways are active in both cancer and normal contexts, it remains unclear how nutrient availability influences the balance between synthesis and salvage.

Extensive study of how glucose, amino acid, and other carbon and nitrogen substrate availability affects nucleotide synthesis and cell proliferation has established that salvage precursors can compensate when synthesis is limiting^6,7,16,17^. Comparatively less attention, however, has been paid to how micronutrients, including vitamins, minerals, and trace elements, impact metabolism under physiologic conditions. This variable is particularly relevant for cancer, where folate-mediated one-carbon metabolism has been implicated as an important regulator of nucleotide synthesis and therapeutic response^18–21^, and where chemotherapies such as methotrexate^22^ directly target vitamin-dependent metabolic pathways. Indeed, anti-folate chemotherapies were found to be effective for treatment of childhood leukemias in part because folate species were recognized as potential drivers of disease progression^23^. Nevertheless, standard cell culture media contain supraphysiologic levels of vitamins and cofactors, including folates^12,20,24,25^. Folates mediate one-carbon transfer reactions required for both purine and thymidine synthesis, while trace metals like iron and zinc regulate enzymes involved in nucleotide metabolism and DNA replication^18,20,26^. Thus, micronutrient abundance or limitation could affect nucleotide metabolism in physiologically relevant contexts. Indeed, physiological nutrient conditions are known to affect how well cancers can grow^12,24,27–29^. In some leukemia contexts, this has been attributed to vitamin levels^30,31^, but the nutrients and micronutrients that limit cancer proliferation for most contexts are not known.

To identify factors that determine how cells acquire nucleotides in physiological conditions to support proliferation, we studied the nucleotide acquisition strategies used by B-cell acute lymphoblastic leukemia (B-ALL). We selected B-ALL as a model because this is a cancer where anti-metabolite therapies targeting nucleotide metabolism are effective, and involve the folate analog methotrexate, which inhibits *de novo* synthesis, and 6-mercaptopurine (6-MP), a thiopurine analog that impairs both synthesis and salvage^22,23^. Furthermore, prior work has shown that combinatorial targeting of *de novo* synthesis, salvage, and DNA replication stress responses can affect nucleotide levels and impair proliferation of B-ALL cells^32,33^. Finally, because blood and bone marrow are physiological environments for B-ALL, and nutrient availability in blood and bone marrow is similar^34^, we can model plasma-like nutrient conditions *in vitro* to dissect mechanisms and confirm the findings in mouse models.

In this study, we first characterized the metabolite composition of plasma from mice with B-ALL and used these measurements to formulate a mouse plasma-like media (MPM) that recapitulates the circulating nutrient environment of leukemic mice. We find that B-ALL cells cultured in MPM acquire nucleotides through both synthesis and salvage pathways, with a specific reliance on thymidine salvage to support dTTP production. Importantly, thymidine salvage involving thymidine kinase 1 (TK1) is required for optimal proliferation in MPM, and TK1 disruption leads to dNTP depletion, DNA replication stress, and impaired cell proliferation. Mechanistically, we find that physiological folate levels are inherently limiting for nucleotide synthesis, constraining *de novo* synthetic flux despite access to physiological levels of glucose and amino acids, and thereby creating a dependence on salvage pathways that cannot be fully compensated by *de novo* synthesis. Consequently, loss of TK1 impairs B-ALL proliferation in blood, bone marrow, and spleen, and proliferation can be modulated by dietary folate availability, demonstrating that endogenous micronutrient constraints shape nucleotide acquisition strategy and can determine the metabolic dependencies of cancer cells *in vivo*.

## Results

### Formulation of mouse plasma medium (MPM) to model how physiological nutrients impact B-ALL metabolism

To model a physiological nutrient environment for B-ALL, we first characterized metabolites in plasma in leukemic and non-leukemic mice. For these studies, we used a well-studied transplantable mouse model of Philadelphia chromosome–positive (Ph^+^) B-ALL established by retroviral expression of p185 BCR-ABL in p19^Arf−/−^ pre-B cells from a C57BL/6J background^35,36^. Following transfer into syngeneic C57BL/6J mice, or immunodeficient NSG mice, disease develops rapidly, with aggressive polyclonal leukemia evident within 2 weeks^35–37^. Quantitative metabolite profiling was performed on plasma isolated from cohorts of 6-8-week-old C57BL/6J and NSG mice approximately 2 weeks after B-ALL cell implantation, a time point corresponding to high leukemic burden, as well as age-matched controls. Disease status was the dominant driver of variance between plasma samples analyzed, with principal component analysis separating leukemic from healthy plasma in C57BL/6J and NSG mice (Fig. 1a; Extended Data Fig. 1a). Of note, metabolite concentrations were highly correlated between C57BL/6J and NSG mice in healthy and leukemic states, indicating that circulating metabolite composition is largely determined by disease state rather than mouse strain or host immune background (Extended Data Fig. 1b,c). Among the most prominent disease-associated differences was elevation of nucleotide species in B-ALL plasma (Fig. 1b; Extended Data Fig. 1d-j). Of note, plasma from patients with Ph^+^ B-ALL also exhibits changes in nucleotide species levels^34^, and cross-species correlation analysis confirmed strong concordance between mouse and human plasma metabolite profiles from both healthy and leukemic contexts (Extended Data Fig. 1k-n). Together, these results identify changes in circulating nucleotide-related species as a conserved feature of Ph^+^ B-ALL in both humans and mice and raise the possibility that this change in nutrient availability may influence how leukemia cells acquire nucleotides to support disease progression.

**Figure 1.**
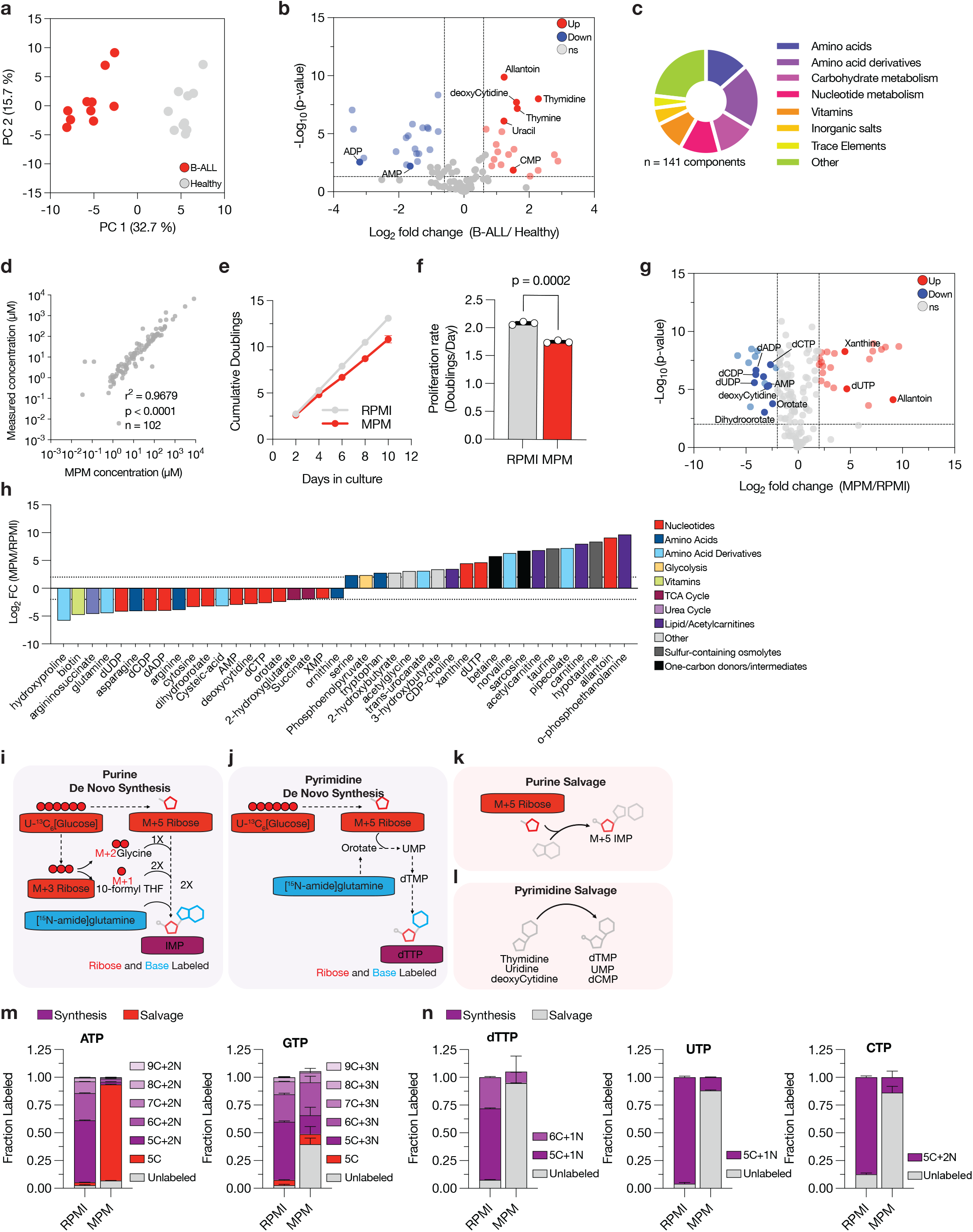
Formulation of mouse plasma-like medium (MPM) to model a physiological nutrient environment for B-cell acute lymphoblastic leukemia (B-ALL). **a**, Principal component analysis (PCA) of metabolite concentrations measured in plasma from healthy and B-ALL–bearing C57BL/6J (BL6) mice. A total of 115 metabolites were measured and contribute to each sample. Data represent n = 10 biologically independent samples per group. **b**, Volcano plot of polar metabolite concentrations in plasma from B-ALL–bearing versus healthy BL6 mice. Metabolites related to nucleotide metabolism that differ between groups are labeled. Significance was defined as |log2 fold change| > 0.6 and raw P < 0.05, n = 10 biologically independent samples per group. **c**, Pie chart showing the fraction of molecules from different classes used to formulate MPM. **d**, Scatter plot of metabolite concentrations measured by LC-MS in reconstituted MPM plotted against the target concentrations used in the MPM formulation. Each data point represents the mean of n = 4 biologically independent measurements. r^2^ and P value were determined by Pearson correlation. **e-f**, Cumulative population doublings of B-ALL cells cultured in RPMI or MPM over time (e) and (f) corresponding proliferation rates (doublings per day) calculated from data presented in (e). Data are mean ± SD, n = 3 biologically independent samples. Significance was determined by unpaired two-tailed t-test. **g-h**, Volcano plot (g) and pathway-level categorization (h) showing log2 fold change in metabolites measured from B-ALL cells cultured in MPM relative to the same cells cultured in RPMI for 8 days. Metabolites related to nucleotide metabolism that differ between groups are labeled. In (g), significance was defined as |log2 fold change| > 2 and raw P < 0.05. In h, the top 20 upregulated and top 20 downregulated metabolites with raw P < 0.05 were selected and annotated by pathway. Horizontal lines in (h) indicate |log2 fold change| = 2. Data represent n = 8 biologically independent samples. **i-l**, Schematic showing *de novo* purine (i) and pyrimidine (j) synthesis pathways and salvage purine (k) and pyrimidine (l) pathways, including the isotope labeling expected from [U-^13^C_6_]glucose and [amide-^15^N]glutamine for each pathway. **m-n**, Fractional labeling of the indicated purine (m) and pyrimidine (n) species measured by LC–MS when cells are exposed to [U-^13^C_6_]glucose and [amide-^15^N]glutamine for 24 hours prior. Data are mean ± SD and represent n = 4 biologically independent samples.

To study how plasma levels of metabolites impact B-ALL cell metabolism to support proliferation, we formulated a mouse plasma-like medium (MPM) based on metabolite concentrations measured in the plasma of leukemic mice. We reasoned that this approach would allow us to most directly compare findings with phenotypes in the transplantable mouse model. In total, MPM was formulated to contain 141 defined components, including polar metabolites, salts and trace elements. Polar metabolite concentrations were derived from B-ALL plasma measurements or from prior studies quantifying metabolites in mouse plasma^9,12^, trace elements were incorporated based on the physiological medium Plasmax^24^, and salts were included at concentrations found in RPMI. 10% dialyzed fetal bovine serum (as a source of lipids and growth factors) and β-mercaptoethanol were included, as in standard B-ALL culture conditions^37–39^ (Fig. 1c; Extended Data Fig. 2). Quantitative LC–MS analysis of the formulated MPM confirmed that metabolite concentrations closely matched those measured in plasma from mice with B-ALL (Fig. 1d). Compared with standard cell culture media, MPM incorporates an expanded set of nutrients to represent the diversity of metabolites detected in plasma in the Human Metabolome Database (HMDB), including nucleotide species and vitamins (Extended Data Fig. 2). Importantly, MPM supported continuous stable proliferation of both murine and human (SUP-B15) Ph^+^ B-ALL cells, as well as K562 cells, although in some cases at a slightly slower rate than that observed in RPMI (Fig. 1e–f; Extended Data Fig. 3a–b). These data confirm that the nutrients found in the plasma of mice with B-ALL, and therefore in MPM, are sufficient to support leukemia cell proliferation.

### B-ALL cells use salvage to obtain nucleotides in plasma-like nutrient conditions

We next asked whether culturing B-ALL cells in MPM changes nucleotide metabolism relative to standard culture medium. Untargeted metabolomics analysis of mouse B-ALL cells cultured in MPM or RPMI revealed distinct intracellular metabolite profiles, with prominent changes observed in levels of nucleotide-related metabolites (Fig. 1g-h; Extended Data Fig. 3c-d). Cells can acquire nucleotides either through *de novo* synthesis or by salvaging nucleotide precursors from the extracellular environment^4^. To quantify the relative contribution of each pathway in B-ALL cells cultured in RPMI or MPM, we traced incorporation of [amide-^15^N]-glutamine and [U-^13^C_6_]-glucose into purine and pyrimidine nucleotides. During *de novo* synthesis, the amide nitrogen of glutamine labels the nucleobase, while glucose-derived carbons contribute to both the ribose and nucleobase moieties (Fig. 1i–l). Thus, nucleotides labeled at both the ribose and nucleobase must arise from *de novo* synthesis; those labeled only in the ribose reflect salvage of an unlabeled base onto a newly synthesized ribose; and fully unlabeled nucleotides arise from direct salvage of intact nucleosides or are pre-existing. That is, nucleotide isotopologues heavier than M+5 correspond to newly synthesized purines or pyrimidines (Fig. 1i-j), while salvage of purine bases by enzymes such as HPRT or APRT yield nucleotides labeled only on the ribose (M+5 isotopologue) (Fig. 1k), and salvage of pyrimidine nucleosides by enzymes such as UCK, dCK, or TK1 yields unlabeled nucleotide species (Fig. 1l). As is inherent to isotope tracing approaches, ribose-only labeled and fully unlabeled species cannot be formally distinguished from pre-existing pool contributions; these are collectively designated as salvage throughout. These labeling patterns therefore enable us to distinguish *de novo* synthesis from salvage, and to assess the relative contributions of each pathway under different nutrient conditions.

After 24 hours of mouse B-ALL cell culture in RPMI with isotope-labeled glucose and glutamine, most purine and pyrimidine nucleotides measured in cells are heavier than M+5, consistent with predominant use of *de novo* synthesis to acquire nucleotides (Fig. 1m-n). This finding is expected, as RPMI lacks nucleotides, nucleosides or nucleobases to support salvage. A small fraction of purine nucleotides in cells cultured in RPMI exhibited ribose-only labeling (Fig. 1m), consistent with either recycling of nucleobases from intracellular nucleotides or a contribution from salvage of trace nucleobases present in the serum added to the media. In contrast, when cells were cultured in MPM, a large fraction of ATP was M+5 labeled, and a large fraction of dTTP, UTP, and CTP were unlabeled, consistent with an increased contribution from salvage (Fig. 1m-n). Despite robust labeling patterns consistent with ATP derived from salvage pathways, GTP labeling suggested a large fraction was derived from *de novo* synthesis in MPM. This pattern is consistent with the composition of MPM, which contains AMP and IMP that can be converted into adenylate nucleotide species^40^, but lacks salvageable guanylate species (Extended Data Fig. 2); although IMP can feed into both adenylate and guanylate pools, the preferential labeling of ATP suggests flux from salvaged IMP is directed toward AMP under these conditions (Fig. 1m). These findings confirm expectations that when salvageable precursors are present, salvage pathways will contribute to nucleotide acquisition, and are consistent with prior studies assessing nucleotide salvage in mouse models^10^. In support of differential use of synthesis and salvage pathways to acquire nucleotides in RPMI compared to MPM, the nucleotide synthesis inhibitors lometrexol, methotrexate, and pemetrexed, which impair one-carbon–dependent thymidylate and purine production, as well as brequinar, which inhibits pyrimidine synthesis, were more effective at inhibiting proliferation of B-ALL cells in RPMI (Extended Data Fig. 3e-h). Conversely, the equilibrative nucleoside transport inhibitor dipyridamole was more effective at inhibiting proliferation in MPM (Extended Data Fig. 3i).

### Nucleotide salvage supports DNA replication and proliferation in plasma-like nutrient conditions

To test whether nucleotide salvage is important for supporting proliferation in plasma-like conditions, we formulated a version of MPM lacking all nucleotides, nucleosides and nucleobases (MPM-Nucleotides). As a control, we also formulated RPMI supplemented with nucleotides and nucleotide precursors found at the same levels as those present in MPM (RPMI+Nucleotides). Addition of nucleotides to RPMI had no effect on B-ALL cell proliferation; however, removing salvageable nucleotides/nucleotide precursors from MPM markedly attenuated proliferation of both mouse and human Ph^+^ B-ALL cells (Fig. 2a,b). These data confirm salvage is important for B-ALL proliferation under plasma-like conditions. Intracellular metabolite profiling revealed that while ribonucleotide levels remained largely unchanged when cells are cultured in MPM with or without nucleotides (Extended Data Fig. 4), dNTPs, most notably dATP and dTTP, were markedly depleted when B-ALL cells are exposed to MPM conditions that lack salvageable precursors, whereas dCTP levels increased (Fig. 2c,d; Extended Data Fig. 4). Such reciprocal shifts align with known allosteric control of ribonucleotide reductase, where changes in levels of specific dNTPs impact substrate specificity^41^.

**Figure 2.**
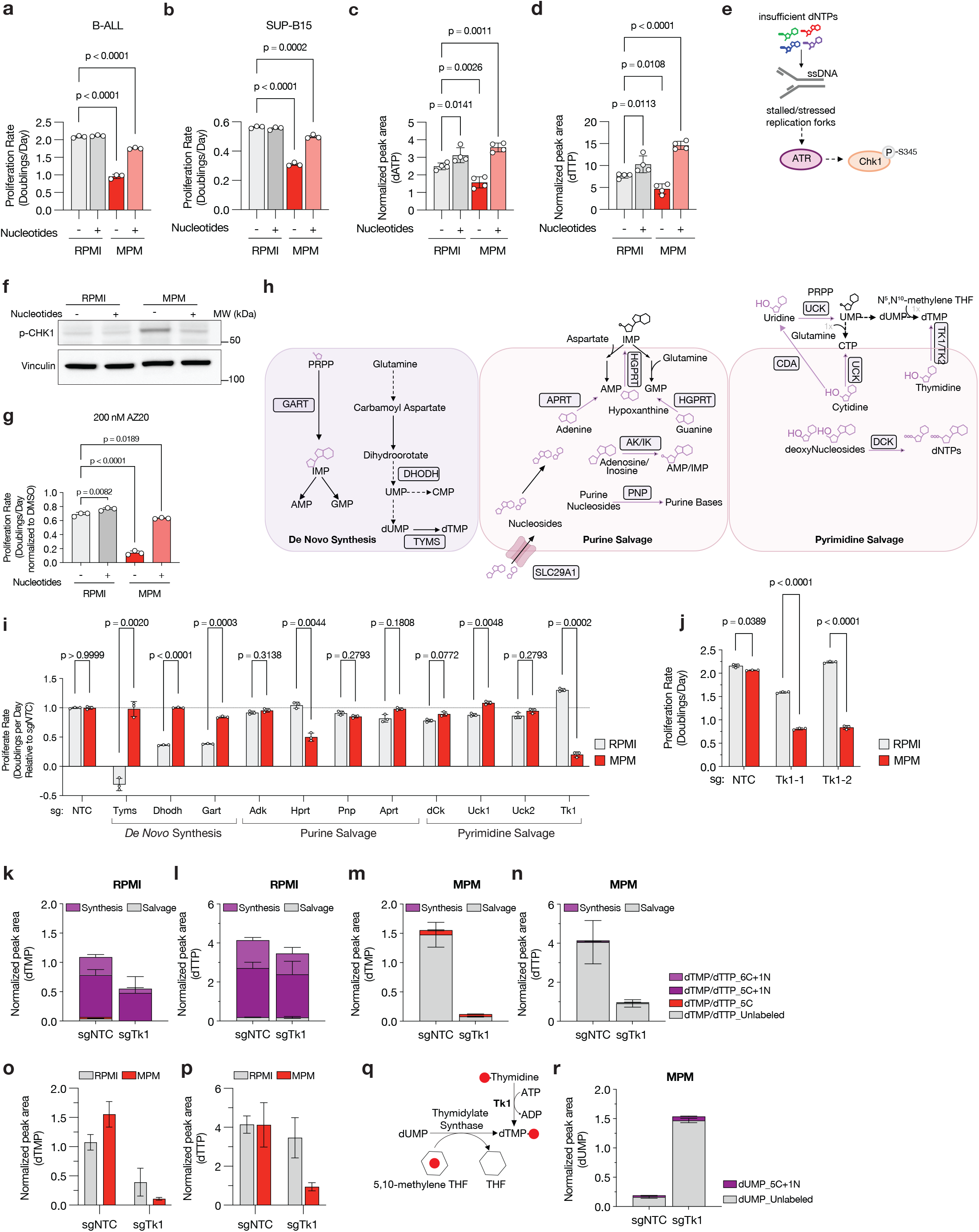
Nucleotide salvage precursors are required for proliferation in MPM nutrient conditions. **a-b**, Proliferation rates (doublings per day) of B-ALL (a) and SUP-B15 (b) cells cultured in RPMI or MPM with or without MPM-level nucleotides for 4 days. Data are mean ± SD and represent n = 3 biologically independent experiments. Statistical significance was determined by ordinary one-way ANOVA followed by Holm–Šídák multiple comparisons test using a pooled variance. **c-d**, LC–MS quantification of intracellular dATP (c) and dTTP (d) levels in B-ALL cells cultured in RPMI or MPM with or without MPM-level nucleotides for 4 days. Data are mean ± SD and represent n = 4 biologically independent samples. Significance was determined by ordinary one-way ANOVA followed by Holm–Šídák multiple-comparisons test. **e**, Schematic showing DNA replication stress signaling in response to dNTP insufficiency. **f**, Western blot analysis of phospho-Chk1 (P-Chk1 S345) in B-ALL cells cultured in RPMI or MPM with or without MPM-level nucleotides for 4 days. Vinculin is shown as a loading control. **g**, Relative proliferation rates (doublings per day, normalized to DMSO control) of B-ALL cells treated with 200 nM AZ20 when cultured in RPMI or MPM with or without MPM-level nucleotides for 4 days as indicated. Data are mean ± SD and represent n = 3 biologically independent experiments. Significance was determined by ordinary one-way ANOVA followed by Holm–Šídák multiple-comparisons test. **h**, Schematic of nucleotide synthesis and salvage pathways highlight a subset of enzymes. **i**, Proliferation rates (doublings per day, normalized to sgNTC) of B-ALL cells cultured in RPMI or MPM following expression of a control guide (sgNTC) or a guide to disrupt the indicated genes involved in nucleotide *de novo* synthesis or salvage pathways. Data are mean ± SD and represent n = 3 biologically independent experiments. Significance was determined by ordinary one-way ANOVA followed by Holm–Šídák multiple-comparisons test. **j**, Proliferation rates (doublings per day) of clonal control (NTC) or TK1 knockout B-ALL cells cultured in RPMI or MPM for 4 days. Data are mean ± SD and represent n = 3 biologically independent samples. Significance was determined by unpaired two-tailed t-tests with Welch’s correction and adjusted for multiple comparisons using the Holm–Šídák method. **k-n**, Isotopologue distributions of dTMP in RPMI (k), dTTP in RPMI (l), dTMP in MPM (m), and dTTP in MPM (n) from B-ALL cells without (sgNTC) or with (sgTK1) TK1 knockout when cells are exposed to [U-^13^C_6_]glucose and [amide-^15^N]glutamine for 24 hours prior. Data are mean ± SD and represent n = 3 biologically independent samples. o-p, Normalized peak areas of dTMP (o) and dTTP (p) from the same experiment in (k-n). **q**, Schematic of thymidine salvage via TK1 or *de novo* dTMP synthesis from dUMP. **r**, Normalized peak areas of dUMP measured in B-ALL cells without (sgNTC) or with (sgTk1) TK1 knockout cultured in MPM exposed to [U-^13^C_6_]glucose and [amide-^15^N]glutamine for 24 hours prior. Data are mean ± SD and represent n = 3 biologically independent samples.

Because imbalances or deficiency of dNTPs can disrupt DNA replication and activate DNA-replication stress signaling that can slow or arrest proliferation^3,33^ (Fig. 2e), we assessed phosphorylation of the kinase Chk1 as a marker of this response. Cells cultured in MPM–Nucleotides exhibited robust Chk1 Ser345 phosphorylation (Fig. 2f), consistent with DNA replication stress^42^. Moreover, cells cultured in MPM-Nucleotides were more sensitive to AZ20, a small molecule inhibitor of ATR, a key kinase involved in the replication stress response, than cells cultured in RPMI or complete MPM (Fig. 2g), suggesting that nucleotide salvage protects cells from DNA replication stress in MPM conditions. Together, these data argue that nucleotide salvage is important to support DNA replication in plasma-like environmental conditions.

To further test the role of nucleotide salvage in supporting proliferation in MPM, we employed CRISPR-Cas9 to individually disrupt several salvage enzymes, including UCK1 and UCK2 (uridine and cytidine salvage), dCK (deoxycytidine salvage), APRT (adenine salvage), HPRT (guanine and hypoxanthine base salvage), PNP (purine nucleoside phosphorylase), and TK1 (thymidine salvage). As a comparison, we also targeted enzymes required for *de novo* nucleotide synthesis, including GART (purine synthesis), DHODH (uridine and downstream cytidine synthesis from UTP) and TYMS (thymidine synthesis) (Fig. 2h). For each gene, we generated polyclonal knockout populations alongside cells expressing a non-targeting control sgRNA (sgNTC). Efficient gene disruption was confirmed by immunoblotting when suitable antibodies were available (Extended Data Fig. 5a-i). For targets where antibodies were not available, to confirm protein-level enzyme loss we instead assessed transcript depletion by qPCR (Extended Data Fig. 5j-k). We then selected guides with the greatest reduction in expression to screen for enzymes that, when lost, impair proliferation in MPM or RPMI. Knockout of nucleotide salvage enzymes had minimal impact on proliferation in RPMI (Fig. 2i). In contrast, loss of the salvage pathway enzymes HPRT and TK1 markedly reduced proliferation in MPM. Conversely, loss of select *de novo* synthesis enzymes impaired proliferation more in RPMI than in MPM. These data are consistent with nucleotide synthesis supporting the majority of nucleotide acquisition in RPMI-based media where salvage precursors are absent, and salvage being a route of nucleotide acquisition in MPM (Fig. 1m-n).

Stable isotope tracing experiments indicated that *de novo* nucleotide synthesis remains active in B-ALL cells cultured in MPM (Fig. 1m-n), suggesting that synthesis still contributes to nucleotide production under plasma-like nutrient conditions. To more directly test the contribution of *de novo* synthesis, and to avoid potential residual activity present in polyclonal knockout populations, we generated clonal B-ALL lines lacking the enzymes DHODH and GART, which are essential for pyrimidine and purine synthesis, respectively. Protein loss was verified by immunoblotting (Extended Data Fig. 6a-b), and functional disruption of *de novo* synthesis was further tested by showing relevant salvage substrates (hypoxanthine and uridine) can rescue the ability of each clone to grow in RPMI (Extended Data Fig. 6c-d). As RPMI does not contain salvage precursors, loss of either enzyme severely impaired proliferation in RPMI (Extended Data Fig. 6e). Notably, however, fully disrupting either enzyme also modestly reduced B-ALL proliferation in MPM, arguing that *de novo* synthesis contributes to nucleotides in MPM even when salvage precursors are present.

Nevertheless, despite the contribution of *de novo* synthesis to proliferation in MPM, CRISPR–Cas9–mediated disruption of nucleotide salvage enzymes revealed that select nucleotide salvage pathways represent the predominant vulnerability under plasma-like conditions. Among the salvage knockouts, TK1 loss resulted in the strongest proliferation defect in B-ALL cells cultured in MPM relative to RPMI, with a phenotype comparable to disruption of DHODH-mediated pyrimidine synthesis in RPMI (Fig. 2i). We therefore focused on thymidine salvage via TK1 to investigate why B-ALL cells are dependent on nucleotide salvage under plasma-like nutrient conditions and generated clonal *Tk1*-knockout (sgTk1) and control (sgNTC) lines. Loss of TK1 protein was confirmed by immunoblotting (Extended Data Fig. 5l) and *Tk1*-deficient clones exhibited a strong proliferation defect in MPM relative to RPMI (Fig. 2j). B-ALL cell clones knocked out for *Hprt* (Extended Data Fig. 5m) similarly proliferated slower in MPM relative to RPMI, and relative to NTC-control clones in MPM (Extended Data Fig. 5n). However, consistent with a more profound proliferation defect in MPM, TK1 loss triggered robust DNA replication-stress signaling, as evidenced by phospho-Chk1 immunoblotting (Extended Data Fig. 5o), suggesting a strong requirement for thymidine salvage to support DNA replication when B-ALL cells are cultured in MPM conditions.

Stable-isotope tracing with [U-^13^C]-glucose and [^15^N-amide]-glutamine confirmed that TK1-deficient cells retain the ability to produce thymidylate nucleotides *de novo* when cultured in RPMI (Fig. 2k-l), but in MPM conditions, extracellular thymidine salvage contributes to dTTP (Fig. 2m-n). Loss of TK1 in cells cultured in MPM led to decreased contribution of extracellular thymidine towards dTMP/dTTP production and also led to reduced dTMP and dTTP levels (Fig. 2m-p), as well as accumulation of dUMP (Fig. 2q-r). These data suggest that the thymidylate synthase step of thymidylate nucleotide synthesis becomes limiting when thymidine salvage is lost. Importantly, TK1 loss did not impair the salvage of other nucleotides in MPM; in fact, levels of ATP, UTP, and dCTP were increased with salvage being the major contributor (Extended Data Fig. 6f-h). These shifts likely are a consequence of ribonucleotide reductase allosteric regulation^41,43^, rather than a direct role for TK1 in non-thymidine metabolism. Nevertheless, TK1-deficient B-ALL cells experience dTTP depletion in MPM nutrient conditions (Fig. 2p).

### Physiological folate concentrations constrain *de novo* nucleotide synthesis in plasma-like nutrient conditions

Because *de novo* synthesis remains active in MPM but cannot sustain nucleotide levels in the absence of salvage precursors (MPM-Nucleotides), we next sought to determine which components of the plasma-like nutrient environment limit this pathway. Compared to RPMI, MPM contains lower concentrations of several precursor metabolites for nucleotide synthesis. To identify which component might be limiting in MPM, we systematically titrated and supplemented media components. Adding increasing amounts of RPMI to MPM-Nucleotides progressively restored proliferation, with as little as 20% RPMI sufficient for restoring proliferation to near RPMI rates (Fig. 3a). Glucose concentrations are comparable between MPM and RPMI, but serine and glutamine levels are lower in MPM (Extended Data Fig. 2), and these amino acids can limit nucleotide synthesis in other contexts^6,44–46^. However, supplementing MPM-Nucleotides with serine and glutamine to RPMI levels had no effect on B-ALL proliferation (Extended Data Fig. 7a). We next asked whether lower folate levels in MPM might constrain proliferation under physiological nutrient conditions. Folates are required cofactors for *de novo* nucleotide synthesis and are included in standard culture media such as RPMI and DMEM at supraphysiological concentrations (2.3–9 µM) compared to both MPM and mouse plasma, where folate species concentrations are ~50 nM (Extended Data Fig. 7b-c) to ~150 nM^47^, both of which are well below the levels in standard media. For MPM formulation, we used 150 nM folic acid (Extended Data Fig. 2; Extended Data Fig. 7c), which falls within this physiological range. Of note, supplementing MPM-Nucleotides with folic acid to match RPMI levels fully restored both mouse and human B-ALL cell proliferation (Fig. 3b; Extended Data Fig. 7d). These data suggest that physiological folate concentrations limit cell proliferation in MPM-Nucleotides.

**Figure 3.**
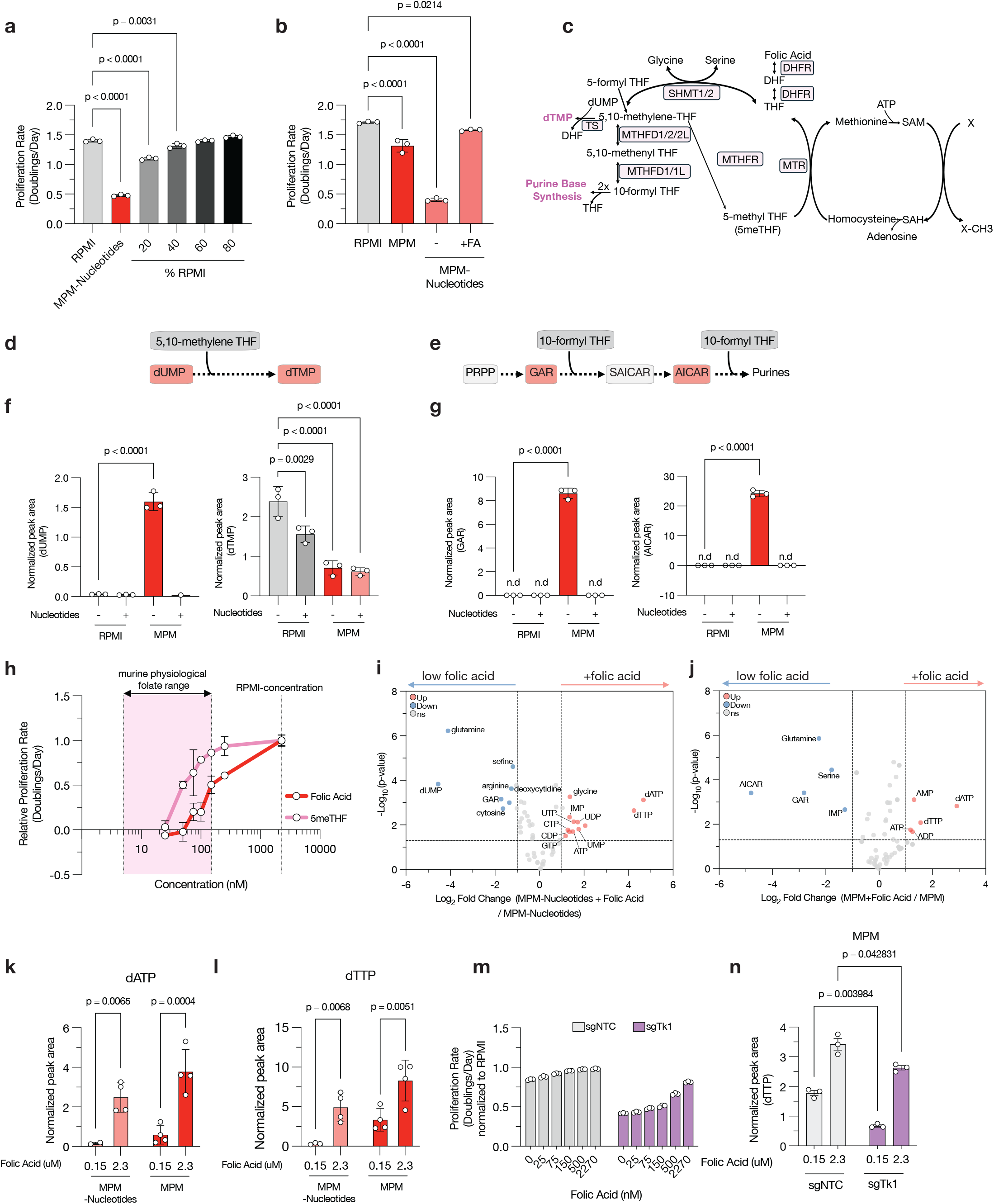
Physiological folate levels limit *de novo* nucleotide synthesis and increase dependence on nucleotide salvage. **a**, Proliferation rates (doublings per day) of B-ALL cells cultured for 4 days in RPMI or MPM without nucleotides with increasing proportions of RPMI as indicated. Data are mean ± SD and represent n = 3 biologically independent experiments. Significance was determined by ordinary one-way ANOVA followed by Holm–Šídák multiple-comparisons test. **b**, Proliferation rates (doublings per day) of B-ALL cells cultured in RPMI, MPM or MPM without nucleotides with or without 2.27 µM folic acid (FA) supplementation. Data are mean ± SD and represent n = 3 biologically independent samples. Significance was determined by ordinary one-way ANOVA followed by Holm–Šídák multiple-comparisons test. **c**, Schematic showing role of folate species in one-carbon metabolism, including contribution to purine and thymidylate synthesis. **d–e**, Schematics showing folate-dependent reactions in dTMP synthesis (d) and in purine synthesis (e). **f–g**, Normalized peak areas measured for the indicated metabolites in B-ALL cells cultured in RPMI or MPM with or without MPM-level nucleotides for 4 days. Data are mean ± SD and represent n = 3 biologically independent samples. Significance was determined by ordinary one-way ANOVA followed by Holm–Šídák multiple-comparisons test. n.d, not detected by LC-MS. **h**, Relative proliferation rates (doublings per day, normalized to 2.27 μM folic acid) of B-ALL cells cultured in MPM without nucleotides containing folic acid or 5-methyl-THF (5meTHF) at the indicated concentrations for 4 days. Data are mean ± SD and represent n = 3 biologically independent experiments. Shaded pink region denotes the physiological range of folate concentrations measured for murine plasma. **i–j**, Volcano plots showing log2 fold changes in metabolite levels measured in B-ALL cells cultured for 4 days in MPM without nucleotides (i) or in MPM (j), with 150 nM folic acid (low folic acid) or 2.27 μM folic acid (+folic acid) as indicated. Significance was defined as |log_2_ fold change| > 1.0 and raw P < 0.05. Data represent n = 4 biologically independent samples per group. **k-l**, Normalized peak areas of dATP (k) and dTTP (l) in B-ALL cells cultured in MPM containing physiological (150 nM) or RPMI-levels of folic acid (2.27 μM). Data are mean ± SD and represent n = 4 biologically independent samples. Significance was determined by two-way ANOVA with interaction followed by Holm–Šídák multiple-comparisons test. **m**, Relative proliferation rates (doublings per day, normalized to proliferation in RPMI) of control (sgNTC) or TK1 knockout (sgTK1) B-ALL cells cultured in MPM formulated with the indicated folic acid concentration. Data are mean ± SD and represent n = 3 biologically independent samples. **n**, Normalized peak areas of dTTP in sgNTC and sgTk1 B-ALL cells cultured in MPM containing physiologic (150 nM) or supraphysiologic (2.27 μM) folic acid. Data are mean ± SD and represent n = 3 biologically independent samples. Significance was determined by unpaired two-tailed Welch’s t-tests with Holm–Šídák correction for multiple comparisons.

Folates serve as carriers for one-carbon units and support multiple metabolic reactions, including methionine and serine metabolism, NADPH generation, and purine and thymidylate synthesis (Fig. 3c). Limitation of folates for nucleotide production could lead to accumulation of pathway intermediates such as dUMP in the thymidylate pathway (Fig. 3d), as well as GAR and AICAR in purine synthesis (Fig. 3e), because the reactions that consume these metabolites require folate-derived one-carbon units. Consistent with this model, cells cultured in MPM-Nucleotides accumulated dUMP (Fig. 3f), as well as GAR and AICAR (Fig. 3g), suggesting that folate limitation constrains nucleotide synthesis in MPM. To further test this idea, we titrated folic acid or 5-Methyltetrahydrofolate (5-methyl-THF), the two primary circulating folate species^48^, across physiological ranges, and increasing levels of either folate species increased proliferation rates (Fig. 3h). Although 5-methyl-THF supported modestly faster proliferation than folic acid when provided individually at the same concentrations, consistent with preferential uptake by the major known folate transporter^49^, physiologically relevant concentrations of either folic acid or 5-methyl-THF were insufficient to sustain maximal proliferation (Fig. 3h). Folic acid was used in experiments because of its greater chemical stability.

To examine how physiological folate levels constrain nucleotide metabolism, we profiled metabolites in B-ALL cells cultured in MPM with or without nucleotides and with or without 2.27 µM folic acid supplementation, a concentration approximately 15-fold higher than that used in formulation of MPM and equivalent to that included in RPMI. Supraphysiologic folate supplementation increased dATP and dTTP levels and reduced accumulation of dUMP and AICAR when cells were cultured in MPM or in MPM-Nucleotides (Fig. 3i–l; Extended Data Fig. 7e–f). These findings suggest that folate availability impacts intracellular nucleotides even when salvageable precursors are present. At physiological folate concentrations, incorporation of isotope-labeled glucose and glutamine into nucleotides through pathways consistent with *de novo* synthesis was reduced across all nucleotide species with pronounced decreases observed for adenylate and guanylate nucleotides, as well as dTTP, which all depend on folate-mediated one-carbon metabolism (Extended Data Fig. 7g–j). Consistent with a folate-dependent bottleneck in nucleotide biosynthesis, the upstream intermediates GAR and dUMP accumulated when cells were cultured in physiological folate concentrations (Extended Data Fig. 7i). In contrast, both isotope labeling and total abundance of UTP and CTP were relatively unchanged by folate concentration (Extended Data Fig. 7j). The altered dNTP levels in physiological folate conditions (Extended Data Fig. 7e, f) were also accompanied by increased Chk1 phosphorylation and γH2AX (Extended Data Fig. 7k).

To directly test whether physiological folate concentrations limit proliferation in standard media, we generated folate-free RPMI^47^ and titrated folic acid or 5-methyl-THF back to defined concentrations. Cells cultured in RPMI displayed impaired proliferation when folates were absent or present at physiological levels (Extended Data Fig. 7l), phenocopying the proliferative defects observed in MPM-Nucleotides and confirming that supraphysiological folate concentrations in standard media mask a folate-dependent limitation of *de novo* nucleotide synthesis. We next asked whether folate supplementation could rescue the reduced proliferation caused by loss of nucleotide salvage in TK1-deficient B-ALL cells cultured in MPM. Increasing folate concentrations enabled increased proliferation of TK1-deficient cells in MPM (Fig. 3m), increased dTTP levels (Fig. 3n), and reduced DNA replication-stress signaling (Extended Data Fig. 7m). These findings suggest that folate limitation impairs dTTP synthesis and increases reliance on TK1-mediated thymidine salvage in B-ALL cells. Taken together, these data indicate that physiological folate concentrations constrain *de novo* thymidylate and purine synthesis, thereby impacting dNTP levels, inducing replication stress, and slowing cell proliferation.

### B-ALL proliferation *in vivo* is supported by TK1 and is modulated by dietary folates

To confirm that the dependency on nucleotide salvage observed in MPM is recapitulated *in vivo*, we performed competitive transplantation assays using fluorescently labeled B-ALL cells with or without TK1 knockout. GFP-labeled NTC cells were mixed at a 1:1 ratio with BFP-labeled NTC or BFP-labeled TK1 knockout cells and injected into the tail veins of recipient mice. After ~14 days, the relative abundance of GFP- and BFP-labeled leukemia cells was assessed in blood, bone marrow, and spleen by flow cytometry (Fig. 4a, Extended Data Fig. 8a). Total leukemic burden, determined by the proportion of mCherry^+^ B-ALL cells in each tissue, was comparable between mice transplanted with BFP-NTC or BFP-TK1-deficient cells, suggesting that mice reached a similar disease burden at morbidity regardless of TK1 status (Extended Data Fig. 8b). However, in competitive transplantation experiments, BFP-NTC cells remained near the initial 1:1 ratio with GFP-NTC cells while BFP-TK1 knockout cells were consistently depleted relative to GFP-NTC cells across all tissues analyzed (Fig. 4b). These findings demonstrate that TK1-mediated thymidine salvage supports B-ALL cell proliferation in mouse tissues.

**Figure 4.**
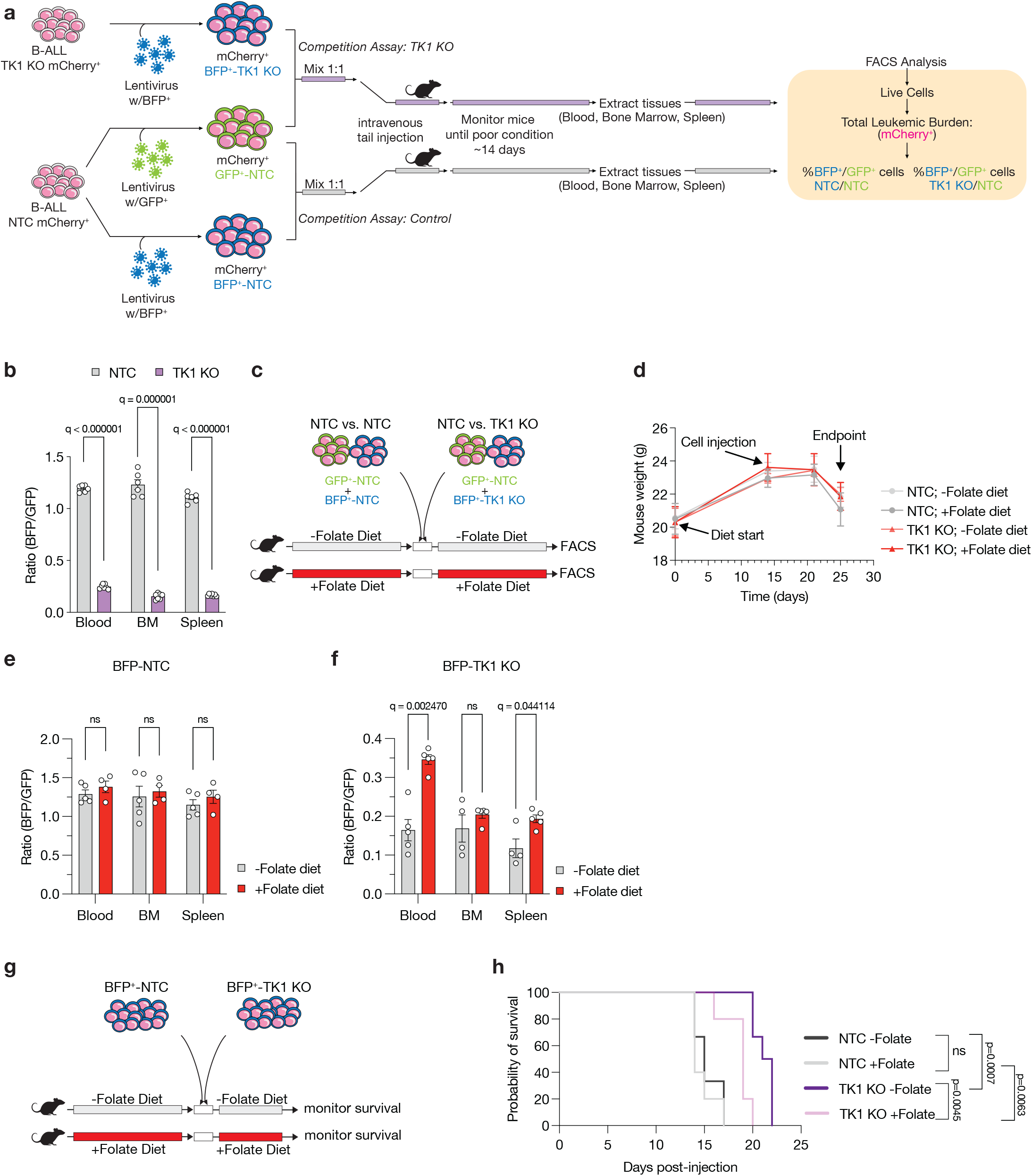
Folate availability influences B-ALL dependency on TK1 *in vivo*. **a**, Schematic depicting competition experiment to assess the relative fitness of B-ALL cells with or without TK1 knockout when implanted into mice. Cas9^+^/mCherry^+^ B-ALL cells were transduced with lentivirus expressing GFP-NTC, BFP-NTC, or BFP-TK1 knockout (KO). GFP–NTC cells were mixed with either BFP-NTC or BFP-TK1 KO cells and injected intravenously into mice. After ~14 days, when mice developed leukemia, blood, bone marrow, and spleen were collected and tumor burden and relative proportions of GFP^+^ and BFP^+^ cells were quantified by flow cytometry. **b**, Ratio of BFP^+^ to GFP^+^ B-ALL cells measured in blood, bone marrow, and spleen from the experiment described in (a). Data are mean ± SD and represent n = 6 biologically independent male NSG mice. Significance was determined by multiple unpaired two-tailed t-tests with Welch’s correction and Benjamini–Krieger–Yekutieli FDR correction. **c**, Schematic depicting competition experiment to assess relative B-ALL fitness (as in (a) with dietary folate modulation. Mice were maintained on folate-depleted (-Folate) or folate-replete (+Folate) diet for 2 weeks before B-ALL transplantation and maintained on the same diet throughout the experiment. **d-f**, Body weight over time of mice maintained on −Folate or +Folate diets (d) and ratio of BFP^+^ to GFP^+^ B-ALL cells in blood, bone marrow, and spleen (e–f) from the competition experiment described in (c). e, BFP-NTC versus GFP-NTC competition. f, BFP-TK1 KO versus GFP-NTC competition. Data are mean ± SEM and represent n = 5 (NTC −Folate; TK1 knockout (KO) −Folate; TK1 KO +Folate) and n = 4 (NTC +Folate) biologically independent female NSG mice. Significance was determined by multiple unpaired two-tailed t-tests with Welch’s correction and Benjamini–Krieger–Yekutieli FDR correction. ns, not significant. **g**, Schematic depicting survival experiment when B-ALL cells without or with TK1 KO are transplanted into mice exposed to diets without or with folates. Mice were maintained on −Folate or +Folate diets for 14 days before implantation of BFP-NTC or BFP-TK1 KO B-ALL cells and remained on the same diets until mice became moribund. **h**, Kaplan–Meier survival curves for mice injected with BFP-NTC or BFP-TK1 KO B-ALL cells under −Folate or +Folate diets as described in (g). Data represent n = 6 (-Folate diet) or n = 5 (+Folate diet) biologically independent NSG male mice per group. Statistical analysis was determined using the log-rank (Mantel–Cox) test.

To determine whether folate availability modulates dependence on TK1 for proliferation *in vivo*, mice were placed on either a folate-free or folate-supplemented diet for two weeks prior to B-ALL cell transplantation and maintained on the same diet during leukemia progression (Fig. 4c). Consistent with previous reports^50^, dietary folate restriction was well tolerated for the duration of the study (Fig. 4d), and plasma levels of folate species were reduced in mice fed the folate-free diet compared to mice fed a folate-supplemented diet (Extended Data Fig. 8c-d). At the experimental endpoint, total leukemic burden was similar across diet groups (Extended Data Fig. 8e-f). However, while folate diet did not affect the relative representation of control competitions (GFP-NTC vs. BFP-NTC) (Fig. 4e), a folate replete diet improved the competitive fitness of TK1 knockout cells relative to control cells (GFP-NTC) (Fig. 4f).

To assess the impact of TK1 loss and dietary folate availability on leukemia progression, mice were maintained on either folate-free or folate-supplemented diets for two weeks prior to implantation of NTC or TK1 knockout B-ALL cells and remained on the same diets throughout the experiment (Fig. 4g). Dietary folate manipulation did not affect survival of mice transplanted with sgNTC cells (Fig. 4h). In contrast, mice transplanted with TK1-deficient cells survived longer than controls, with a greater survival benefit observed in animals maintained on the folate-free diet (Fig. 4h). Together, these findings demonstrate that thymidine salvage via TK1 supports B-ALL proliferation *in vivo* and that dietary folate availability also can impact dependence on nucleotide salvage.

## Discussion

Cancer metabolism research has largely been conducted under nutrient conditions that poorly reflect the physiological environment tumors encounter *in vivo*. Our findings demonstrate that folate availability, a micronutrient routinely oversupplied in standard culture media, constrains *de novo* nucleotide synthesis under physiological conditions. This limitation shifts the balance between synthesis and salvage pathways, increases reliance on salvage, and modulates proliferation in ways that are masked under conventional experimental conditions. Specifically, in B-ALL cells, this dependence on salvage was particularly prominent for TK1-mediated thymidine salvage, consistent with the requirement for folate-derived one-carbon units for thymidylate synthesis. Although not the primary focus of this study, reduced dTTP availability following thymidine salvage loss likely propagates broader dNTP imbalance through allosteric regulation of ribonucleotide reductase^43^. Consistent with this model, TK1 loss decreased dATP and increased dCTP levels, suggesting that TK1 deficiency induces DNA replication stress not only by depleting dTTP, but also by promoting dNTP imbalance.

Although folate-derived one carbon units are also required for purine synthesis, disrupting purine salvage was less of a dependency for the B-ALL cells tested in physiological folate conditions. This could be because multiple purine salvage pathways are active in B-ALL cells, which could enable salvage of purine precursors such as inosine, AMP, and IMP which are all present in MPM at concentrations between 7– 20 µM (Extended Data Fig. 2). Of note, HPRT loss in B-ALL had minimal effect on causing DNA replication stress, possibly because of compensatory purine salvage via adenosine kinase (ADK) and purine nucleoside phosphorylase (PNP), which are expressed in lymphoid tissues and can salvage purines through alternate routes^51^. Prior work has found different cell types exhibit different capacities to salvage various nucleotide precursors from their environment^3,33^, and thus the specific salvage enzymes different cell types are dependent on may vary. Salvageable nucleotide precursor availability and cancer type will also contribute to which nucleotides are most affected by physiological folate-induced constraints on nucleotide synthesis. Because nucleotide and nucleotide precursor levels vary widely across tissues^9,11^, and the ability to access those metabolites is affected by differential expression of salvage enzymes^3,33,52^, how constraints on nucleotide synthesis impact reliance of different cancers on specific salvage precursors is expected to be context dependent.

Species-specific differences in nucleotide metabolism may also play a role in how physiological folate-driven constraints on nucleotide synthesis affect nucleotide levels in cells. For example, in humans, thymidine phosphorylase (TYMP) is broadly expressed and converts thymidine to thymine and deoxyribose-1-phosphate^53^, potentially reducing circulating thymidine levels and making blood cells more reliant on other salvageable pyrimidine species. While we observed elevated plasma nucleotides in both B-ALL–bearing mice and humans^34^, the specific nucleotide species elevated differ between species. These distinctions emphasize that the salvageable nucleotide precursors available in blood is shaped by both enzymatic activity and species-specific plasma composition, which may in turn influence which salvage enzymes are essential for cell proliferation.

These findings underscore the importance of considering dietary influences on circulating nutrients when modeling cancer metabolism. Standard breeding chow used in many research animal facilities contains high levels of folic acid, often exceeding 2 mg/kg, which can elevate plasma folate levels beyond physiological norms and mask folate-dependent metabolic vulnerabilities^54^. This has implications for preclinical assessment of cancer therapies, especially those that might impact nucleotide metabolism. It may also contribute to why mouse models are sometimes poor predictors of anti-metabolite chemotherapy efficacy in patients^55,56^.

CRISPR screening datasets, such as DepMap performed under nutrient-replete standard culture conditions, identify strong dependencies on *de novo* nucleotide synthesis genes in hematologic malignancies^33,57^ (Extended Data Fig. 9), while suggesting tolerance to nucleotide salvage gene loss. These findings likely reflect screening performed under nutrient-replete standard cell culture conditions and suggest that metabolic dependencies identified *in vitro* incompletely capture those operating in physiological environments. Screening approaches that better model circulating nutrient availability may therefore reveal context-dependent vulnerabilities that are masked under conventional culture conditions^11,58^. Additionally, redundancy between *de novo* synthesis and salvage pathways may buffer proliferating cells against single-agent metabolic inhibition. Prior studies demonstrate that simultaneous disruption of complementary nucleotide pathways can enhance replication stress^32,59,60^, emphasizing the potential of targeting multiple arms of nucleotide metabolism to exploit vulnerabilities in nutrient-restricted microenvironments.

More broadly, these findings underscore the importance of modeling micronutrient availability when studying cancer metabolism. Work using various physiological media formulations, including HPLM^12^, Plasmax^24,61^ and serum-derived media^58^, has demonstrated how macronutrient environment can impact metabolism; however, the impact of micronutrients have been less studied and are not always modeled well even by physiological media formulations. Folates represent one such example^31^, but similar discrepancies exist for other essential cofactors, including riboflavin^30^, thiamine, and metal ions such as iron^62^, zinc and copper, that influence one-carbon transfer, redox balance, and DNA synthesis^30^. These metabolites circulate at nanomolar to micromolar levels yet are often supplied in culture at much higher concentrations. Overall, our findings show that systematically defining how the availability of salvage precursors, vitamins, and biosynthetic substrates interact to shape nucleotide metabolism will provide clearer insight into how rapidly proliferating cells meet nucleotide demand. More broadly, accurately modeling these nutrient interactions in physiologically relevant conditions will be essential for understanding variable responses to therapies targeting nucleotide metabolism.

## Lead Contact

Further information and requests for resources and reagents should be directed to and will be fulfilled by Matthew G. Vander Heiden.

## Acknowledgements

We thank members of the Vander Heiden laboratories for helpful discussions and the Koch Institute’s Robert A. Swanson (1969) Biotechnology Center for technical support, specifically the Flow Cytometry facility. We also thank the MIT Division of Comparative Medicine staff for help with colony maintenance and animal care. We thank Wontaek Chung and Paul Chamberlin for intravenous tail injections. We thank the Stegmaier Laboratory (Dana-Farber Cancer Institute) for providing SUP-B15 cells. This work was supported in part by the Koch Institute Cancer Center Support Grant P30CA014051. R.E. was supported by an HHMI Gilliam Fellowship. K.L.A. was supported by the National Science Foundation (DGE-1122374) and National Institutes of Health (NIH) (F31CA271787, T32GM007287). A.A. received support as a Howard Hughes Medical Institute (HHMI) Medical Research Fellow. YG was supported by a Ludwig Cancer Center at MIT Postdoctoral Fellowship. S.S. acknowledges support from the Damon Runyon Cancer Research Foundation (DRG-2367-19). B.D. was supported by National Heart, Lung, and Blood Institute grant F30HL156404. M.G.V.H. acknowledges support from the MIT Center for Precision Cancer Medicine, the Ludwig Center at MIT, and the NIH (R35CA242379).

## Author contributions

Conceptualization: R.E., K.L.A., A.A., M.G.V.H.; Methodology: R.E., K.L.A., A.A., M.G.V.H.; Investigation: R.E., K.L.A., D.L.R., A.A., A.S.Z., A.M.B., M.W., A.W., Y.G., B.T.D., S.S., A.R., T.K., M.W., E.H.R., M.B.M.,; Resources M.T.H., M.G.V.H.; Writing – Original Draft: R.E., K.L.A., M.G.V.H.; Writing – Review & Editing: All authors; Supervision: M.T.H., M.G.V.H.; Funding Acquisition: M.G.V.H.

## Materials and Methods

### Cell lines and standard cell culture

Murine BCR–ABL1^+^; p19^Arf-/-^; mCherry^+^ B-ALL cells^35,36^ were maintained in RPMI-1640 (Corning Life Sciences, 10-040-CV) supplemented with 10% heat-inactivated fetal bovine serum (FBS; Gibco, 10437-028 for passaging media, F0392, Sigma-Aldrich for experimental media) and 50 µM β-mercaptoethanol (Gibco, 21985023). Cas9^+^; BCR–ABL1^+^; p19^Arf-/-^; mCherry^+^ B-ALL cells^37^, were cultured in the same maintenance medium with the addition of 10 µg/mL blasticidin. K562 cells were maintained in RPMI-1640 supplemented with 10% heat-inactivated FBS and 50 µM β-mercaptoethanol. SUP-B15 human B-ALL cells were cultured in RPMI-1640 supplemented with 4 mM L-glutamine, 50 µM β-mercaptoethanol, and 20% heat-inactivated FBS. Lenti-X 293T cells (Takara Bio, 632180) were maintained in DMEM (Corning Life Sciences, 10-017-CV) supplemented with 10% heat-inactivated FBS. All cell lines were routinely tested for mycoplasma contamination using the MycoAlert PLUS Mycoplasma Detection Kit (Lonza BioSciences, LT07-710) and cultured at 37 °C with 5% CO_2_.

RPMI 1640 media (15-040-CV, Corning) lacking folic acid was prepared as described previously^28^. Briefly, RPMI 1640 powder was formulated from individual components to generate 25 L of medium by creating two separate pools: one containing amino acids, inorganic salts, glutathione, and phenol red, and a second containing vitamins excluding folate. The two mixtures were homogenized using an electric blade coffee grinder (Hamilton Beach, 80365) that had been washed three times with methanol and rinsed with distilled water. Homogenized powders were stored at −20 °C until use. For media preparation, powders were weighed and combined and dissolved in water prior to sterile-filtration through a 0.22-µm membrane. To supplement folates, folic acid (F8758, Sigma-Aldrich) or (6S)-5-methyl-5,6,7,8-tetrahydrofolic acid (5-methyl THF) (Schircks Laboratories, 16.236) were dissolved in water to make a stock solution of >1 mM and, immediately prior to each experiment, working stocks were prepared at 100 µM to supplement media to the desired final folate concentration. Hypoxanthine, uridine, and thymidine stock solutions (100 mM) were prepared and stored at −20 °C. Hypoxanthine was dissolved in 1 N NaOH, whereas uridine and thymidine were dissolved in water.

### Formulation of Mouse Plasma Medium (MPM)

MPM was formulated to match the concentrations of metabolites measured in plasma from leukemic B-ALL mice, with some additional metabolites and trace elements incorporated based on prior physiological media formulations^24^ or reported concentrations in mouse plasma^9,12^. The composition of each metabolite pool and the identities and concentrations of all components are listed in Supplementary Table 1. MPM was assembled from seven metabolite pools. Six pools were generated by weighing dry metabolite powders, dissolving them in water, and freezing the solutions at −20 °C as single-use aliquots. The seventh pool, containing salts, was prepared by weighing components and homogenizing them using an electric blade grinder (Hamilton Beach, 80365) washed with methanol and water. The seven pools were combined, brought to volume with water, and supplemented with 10% dialyzed FBS prior to sterile-filtration through a 0.22-µm membrane. To modify levels of metabolites in MPM, such as MPM-Nucleotides, media was formulated by omitting the nucleotide/nucleoside pool and reconstituting the remaining six pools. Amino-acid–supplemented MPM variants were produced by modifying the amino acid pool to supplement serine and glutamine to RPMI-equivalent concentrations. Folate-titrated MPM was generated by preparing MPM metabolite pools lacking folate species (5-methyl-THF and folic acid) and adjusting the concentration of folic acid or 5-methyl-THF as above for standard media.

For B-ALL, SUP-B15, and K562 cell culture, media were further supplemented with 50 µM β-mercaptoethanol.

### Cell Proliferation and Viability

B-ALL cells were pre-cultured in the indicated media (RPMI, MPM or modified MPM formulations without folates to deplete intracellular folates^47^ for 48 hours prior to plating). For plating or passaging, live cell counts were obtained by mixing cells with an equal volume of 0.2% trypan blue and measuring them using a Cellometer Auto T4 Brightfield Cell Counter (Revvity, CMT-AT4P). Cells were seeded for experiments at 250,000 cells per mL, with treatments applied as indicated in the figure legends. For initial day-0 cell counts in proliferation assays and for normalization of intracellular metabolite levels, cell numbers were measured using a Multisizer 3 Coulter Counter (Beckman Coulter) with a diameter range of 7.5–30 µm. Proliferation rates (doublings per day), were calculated as log_2_(final cell count/initial cell count) divided by the number of days.

To assess viability following drug treatment, B-ALL cells were pre-cultured for 48 hours in the indicated medium prior to treatment. Cells were seeded at 250,000 cells per mL in 2 mL of media in 6-well plates, exposed to drug at the indicated concentration, and viability was assessed after 48 hours by live cell counts using a Multisizer 3 Coulter Counter (Beckman Coulter). Viability was normalized to DMSO vehicle-treated controls. The following compounds and vendors were used: Methotrexate (Sigma-Aldrich, A6770-10MG), brequinar (Cayman Chemical Company, 24445), pemetrexed (Selleck Chemicals, S1135), lometrexol (Cayman Chemical Company, 18049), dipyridamole (Tocris, 0691) and the ATR inhibitor AZ20 (Selleck Chemicals, S7050).

### Genetic manipulation of cells

Lenti-X 293T cells were transfected with the lentiviral packaging plasmids pMD2.G (Addgene, 12259), pMDLg/pRRE (Addgene, 12251), and pRSV-Rev (Addgene, 12253) using TransIT®-293 Transfection Reagent (Mirus, MIR 2700). Forty-eight hours post-transfection, medium was collected and filtered through a 0.45 µm low–protein-binding membrane. Viral supernatants were used immediately or stored at −80 °C. Cell lines were transduced in the presence of 10 µg/mL polybrene and spinfected at 900 × g for 1.5 h at 37 °C. Following spinfection, cells were incubated for the remainder of the 24 hour transduction period before replacement with fresh medium.

To generate CRISPR/Cas9-mediated knockout cells, polyclonal knockout populations were generated by transducing Cas9^+^ B-ALL cells with lentivirus derived from pRDACrimson_170 (Crimson-Puro-gRNA) encoding sgRNAs targeting the gene of interest or a non-targeting control (NTC) sgRNA. sgRNA sequences were designed by CRISPick (https://portals.broadinstitute.org/gppx/crispick/public), and sequences used in this study are listed in Supplementary Table 3. Twenty-four hours after transduction, cells were selected with 2 µg/mL puromycin for 48 h, and knockout efficiency was confirmed by western blotting or qPCR where antibodies were not available. Clonal *Tk1, Hprt, Dhodh*, and *Gart* knockout lines were generated using the same approach followed by single-cell sorting of Crimson^+^ cells into 96-well plates via FACS. Clonal populations were expanded, and knockout was confirmed via western blotting. The sgRNAs used to generate clonal knockouts were: Tk1, TAGGACTGACCGATCATGTG; Hprt, AACAAATCTAGGTCATAACC; Dhodh: TTGATCCAGAGTCGGCGCAC; Gart: ATGGCGAGTAAAGGGTACCC.

### Western blot

Cells were pelleted by centrifugation, washed once in PBS, and lysed in RIPA buffer containing protease and phosphatase inhibitors (Cell Signaling Technology, 5871). Lysates were rocked for 15 min at 4 °C, and insoluble material was removed by centrifugation at 21,000 × g for 10 min at 4 °C. Protein concentration was quantified by Bradford assay, and samples were mixed with LDS sample buffer (Thermo Fisher Scientific, NP0008) and 2.5% 2-mercaptoethanol and incubated at 95 °C for 5 min. Equal protein amounts were resolved by SDS–PAGE and transferred onto nitrocellulose membranes using a wet-tank transfer system (Bio-Rad). Membranes were blocked in 5% milk in TBST and incubated overnight at 4 °C with primary antibodies diluted in 5% BSA in TBST. After washing, membranes were incubated with HRP-linked anti-rabbit (Cell Signaling Technology, 7074; 1:5,000) or anti-mouse (Cell Signaling Technology, 7076; 1:5,000) secondary antibodies diluted in 5% BSA in TBST, and signal was detected using a digital chemiluminescence imager (GE Healthcare, LAS 4000).

Primary antibodies used were: TK1 (Proteintech, 15691-1-AP, 1:1,000), HPRT1 (Proteintech, 15059-1-AP, 1:2,000), dCK (Thermo Scientific PA5-27787, 1:1,000), PNP (Proteintech 18009-1-AP, 1:1,000), APRT (Thermo Scientific PA5-95613, 1:1,000), TYMS (Proteintech 15047-1-AP, 1:1,000), UCK1 (Thermo Fisher Scientific, 12271-1-AP, 1:1,000), GART (Proteintech, 67939-1-Ig, 1:1,000), DHODH (Santa Cruz Biotechnology sc-166348, 1:1,000), phospho-CHK1 S345 (Cell Signaling Technology 2341S, 1:1,000), Phospho-Histone H2A.X Ser139 (Cell Signaling Technology 20E3, 1:1,000), Vinculin (Cell Signaling Technology 13901S, 1:5,000). When re-probing was required, membranes were incubated in 30% hydrogen peroxide (Sigma-Aldrich, H1009) for 30 minutes at 37 °C to inactivate HRP.

### Flow cytometry

Flow cytometry analysis was performed on BD FACSCanto, LSR II, and Celesta machines. Sorting was performed on a BD FACSAria II using a 70-µm nozzle at 70 psi. Gating and downstream analysis were performed using FACSDiva and FlowJo.

### Mouse studies

All animal experiments were approved by the MIT Committee on Animal Care. C57BL/6J and NOD-SCID-gamma (NSG) male and female mice 6-10 weeks of age were used. Animals were housed at ambient temperature and humidity (18–23 ºC, 40–60% humidity) on a 12 hr light and 12 hr dark cycle and co-housed with littermates with ad libitum access to food and water. All experimental groups were age-matched, numbered, and randomly assigned to treatment or control groups to minimize bias. Randomization was performed using a randomized numbering sequence at the time of injection or treatment. Investigators were not blinded during animal injections or treatments; however, researchers were blinded during data acquisition and quantification. No statistical methods were used to predetermine sample size. Data were collected from distinct animals, where n represents biologically independent samples. Mice were euthanized when humane endpoint criteria (e.g., more than 20% weight loss, body condition score of less than 2 or signs of distress) were observed. These institutional limits were not exceeded in any experiment.

For leukemia transplantation studies, B-ALL cells were resuspended in 100 µL PBS and injected into recipient mice via tail vein. For competition experiments, 100,000 cells were transplanted, and mice were monitored until they developed signs of advanced leukemia (e.g., lethargy, weight loss, or hind-limb paralysis), at which point tissues were collected for analysis. For survival studies, 10,000 cells were transplanted and mice were monitored until they reached humane endpoints consistent with advanced leukemia and were euthanized. For plasma metabolomics experiments, plasma was obtained from leukemia-bearing mice and from PBS-injected control mice harvested in parallel when the leukemia-bearing cohort developed advanced leukemia. Blood was collected by cardiac puncture or submandibular vein bleed into EDTA-coated tubes (Sarstedt, 41.1395.105), centrifuged at 800 × g for 10 min at 4 °C, and the resulting plasma was flash-frozen in liquid nitrogen and stored at −80 °C until analysis.

### Folate supplementation and dietary manipulation

For dietary folate modulation, mice were fed either a folate-deficient diet containing 1% succinylsulfathiazole (SST) and elevated B-vitamin levels (Inotiv, TD.130451) or a folate-control diet containing 1% SST and 0.003 g/kg folic acid (Inotiv, TD.10399). Both diets were purified and irradiated. Mice were maintained on the assigned diet for two weeks prior to leukemia cell transplantation and remained on the same diet for the duration of the experiments.

### *In vivo* competition experiments

For labeled B-ALL cells competition experiments, clonal NTC or *Tk1*-knockout B-ALL cells were transduced with lentivirus derived from pU6-sgRNA EF1α-Puro-T2A-GFP (Addgene, 111596) or pU6-sgRNA EF1α-Puro-T2A-BFP (Addgene, 60955). These constructs did not contain sgRNAs. Forty-eight hours after transduction, cells were sorted by FACS to enrich for GFP^+^ or BFP^+^ populations, followed by a second sort to increase purity. GFP-NTC cells were mixed 1:1 with either BFP-NTC or BFP-*Tk1* knockout cells (total of 100,000 cells), and the starting representation of each population was confirmed by FACS. Gating strategy is provided in Extended Data Fig. 8a. The mixed suspension was injected into recipient mice via tail vein. After 12–14 days, when mice became moribund, blood and tissues were harvested for analysis. Blood was collected by submandibular bleed into tubes containing ACK lysing buffer (Gibco A1049201), incubated for 15 min at room temperature, centrifuged, and resuspended in FACS buffer (2% FBS in PBS). Spleen and bone marrow were processed by gently grinding tissues in 1 mL ACK lysing buffer with a mortar and pestle, followed by addition of 4 mL buffer. Samples were incubated for 5 min at room temperature, filtered through a 40-µm mesh, centrifuged, and resuspended in FACS buffer for flow cytometry.

### Absolute metabolite quantification by mass spectrometry

Metabolite quantification in plasma samples was performed as described previously^9^. In brief, 5 µL of sample or external chemical standard pool (ranging from ~5 mM to ~1 µM) was mixed with 45 µL of acetonitrile:methanol:formic acid (75:25:0.1) extraction mix including isotopically labeled internal standards. All solvents used in the extraction mix were HPLC grade. Samples were vortexed for 15 min at 4 °C and insoluble material was sedimented by centrifugation at 16,000 x g for 10 min at 4 °C. 20 µL of the soluble polar metabolite extract was transferred to LC-MS vials for analysis.

LC-MS analysis was conducted on a QExactive benchtop orbitrap mass spectrometer equipped with an Ion Max source and a HESI II probe, which was coupled to a Dionex UltiMate 3000 HPLC system (Thermo Fisher Scientific). External mass calibration was performed using the standard calibration mixture every 7 days. An additional custom mass calibration was performed weekly alongside standard mass calibrations to calibrate the lower end of the spectrum (m/z 70-1050 positive mode and m/z 60-900 negative mode) using the standard calibration mixtures spiked with glycine (positive mode) and aspartate (negative mode). 2 µL of each sample were injected onto a SeQuant® ZIC®-pHILIC 150 × 2.1 mm analytical column equipped with a 2.1 × 20 mm guard column (both 5 mm particle size; EMD Millipore). Buffer A was 20 mM ammonium carbonate, 0.1% ammonium hydroxide; Buffer B was acetonitrile. The column oven and autosampler tray were held at 25 °C and 4 °C, respectively. The chromatographic gradient was run at a flow rate of 0.150 mL min^-1^ as follows: 0-20 min: linear gradient from 80-20% B; 20-20.5 min: linear gradient from 20-80% B; 20.5-28 min: hold at 80% B. The mass spectrometer was operated in full-scan, polarity-switching mode, with the spray voltage set to 3.0 kV, the heated capillary held at 275 °C, and the HESI probe held at 350 °C. The sheath gas flow was set to 40 units, the auxiliary gas flow was set to 15 units, and the sweep gas flow was set to 1 unit. MS data acquisition was performed in a range of m/z = 70–1000, with the resolution set at 70,000, the AGC target at 1 × 10^6^, and the maximum injection time at 20 ms.

Metabolite identification was performed using XCalibur 2.2 software (Thermo Fisher Scientific) by matching detected features to external standard pools based on m/z and retention time, with a mass accuracy tolerance of 5 ppm and a retention time window of ±0.5 min. The list of metabolites, internal standards, and calibration details follows our previously published workflow^9^, with the analytes quantified in this study listed in Supplementary Table 1.

For metabolites with isotopically labeled standards, peak areas of labeled standards were first compared with external standard dilutions of known concentration to determine the absolute concentration of each internal standard in the extraction solvent. The peak area of each analyte in plasma samples was then compared with the quantified internal standard to determine absolute metabolite concentrations. Metabolites lacking isotopically labeled internal standards were quantified by external calibration. Peak areas were normalized to a labeled amino acid internal standard eluting at a similar retention time. The same normalization was applied to external standard pool dilutions, and standard curves relating normalized peak area to analyte concentration were generated. Only metabolites with linear calibration curves (typically r^2^ ≥ 0.95) were retained for further analysis.

### Metabolite quantification in cells

B-ALL cells growing in culture were collected by centrifugation, washed once with ice-cold blood bank saline, and metabolites were extracted by resuspending cell pellets in 500 µL ice-cold HPLC-grade methanol:water (80:20) containing 500 nM ^13^C/^15^N-labeled amino acids (Cambridge Isotope Laboratories, MSK-CAA-1) as internal standards. Samples were vortexed for 10 min at 4 °C and centrifuged at 16,000 × g for 10 min at 4 °C. The soluble polar metabolite fraction was collected, dried under nitrogen gas, and stored at −80 °C until analysis. Immediately prior to LC–MS, extracts were resuspended in 20 µL water and transferred to LC-MS vials.

LC–MS analysis was performed under the chromatographic and mass spectrometry conditions described above. For targeted analysis of nucleotide species, the same chromatographic conditions were used with the addition of a narrow-range full scan (m/z 220–700) acquired in negative ion mode to enhance nucleotide detection. Relative metabolite abundances were quantified using XCalibur QuanBrowser 2.2 (Thermo Fisher Scientific) with a 5 ppm mass tolerance and referencing an in-house retention time library of chemical standards. Measurements were normalized to the ^13^C/^15^N-labeled valine internal standard and to cell number.

### Quantification of folates in plasma

For plasma folate analysis, 10 µL of mouse plasma was mixed with 90 µL of extraction buffer (80:20 methanol:water supplemented with 2.5 mM sodium ascorbate, 25 mM ammonium acetate, and 100 nM aminopterin). Samples were vortexed for 10 min at 4 °C and centrifuged at 21,000 × g for 10 min at 4 °C. The supernatant was collected and dried under nitrogen gas.

Folate species were analyzed using the same LC–MS instrumentation described above and following previously established methods^47^. Dried extracts were resuspended in 30 µL water, and 15 µL was injected onto an Ascentis Express C18 column (2.7 µm × 15 cm × 2.1 mm; Sigma-Aldrich). The column oven and autosampler tray were maintained at 30 °C and 4 °C, respectively. Chromatographic separation was achieved using buffer A (0.1% formic acid in water) and buffer B (acetonitrile with 0.1% formic acid) at a flow rate of 0.25 mL min^-1^ with the following gradient: 0–5 min, 5% B; 5–10 min, linear increase from 5% to 36% B; 10.1–14.0 min, linear increase from 36% to 95% B; 14.1–18.0 min, return to 5% B. The mass spectrometer was operated in full-scan, positive-ion mode. MS data were acquired using three narrow-range scans (438–450 m/z, 452–462 m/z, and 470–478 m/z) with a resolution of 70,000, an AGC target of 1 × 10^6^, and a maximum injection time of 150 ms. Quantification was performed using Xcalibur QuanBrowser v2.2 (Thermo Fisher Scientific) with a 5-ppm mass tolerance. Folate species were quantified by normalization to the internal standard aminopterin followed by comparison with calibration curves generated from folate chemical standards to determine absolute concentrations.

### Labeled metabolite tracing

B-ALL cells were pre-cultured for 48 hours in the indicated medium (RPMI, MPM, or nutrient-modified MPM formulations) prior to isotope tracing. For labeling experiments, cells were washed once with blood bank saline and resuspended at 500,000 cells per mL in fresh medium in which glucose and glutamine were replaced with [U-^13^C_6_]-glucose (110187-42-3, Cambridge Isotope Laboratories) and [amide-^15^N]-glutamine (59681-32-2, Cambridge Isotope Laboratories) at the same concentrations as their unlabeled counterparts. Cells were cultured under these conditions for 24 hours, unless otherwise indicated. Following the labeling period, cells were collected and metabolites were extracted and analyzed by LC-MS as described above.

### DepMap CRISPR gene dependency analysis

CRISPR gene dependency scores (Chronos algorithm, DepMap release 25Q3+) were obtained from the Cancer Dependency Map (DepMap) portal (https://depmap.org/portal). Chronos scores represent gene effect estimates corrected for copy number and guide efficiency, with more negative values indicating stronger dependency. Leukemia cell lines were extracted and stratified by lineage annotation into lymphoid/myeloid versus non-lymphoid/myeloid groups based on DepMap lineage metadata. Gene sets corresponding to *de novo* nucleotide synthesis and nucleotide salvage pathways were curated based on established pathway annotations and prior literature. For each gene, dependency scores were compared between lineage groups using an unpaired two-sided Mann–Whitney U test. Statistical analyses were performed in Prism and p values < 0.05 were considered statistically significant unless otherwise indicated.

### Statistics and reproducibility

All *in vitro* experiments were independently repeated at least twice with similar results. Mouse experiments were performed once for each condition unless otherwise specified. All n values represent biologically independent samples. Sample sizes, reproducibility and statistical tests used for each figure are denoted in the figure legends or Source Data. All graphs were generated using GraphPad Prism 11.0.0 (GraphPad Software). Relative fold changes were determined by normalizing every data point to the mean value of the control group.

After determining the absolute or relative concentration of each metabolite in each plasma sample, all multivariate statistical analysis on the data was performed using Metaboanalyst (v6.0)^63^. Metabolite concentrations were auto-scaled and metabolites that contained greater than 50% missing values were removed prior to analysis. For the remainder of the metabolites, missing values were replaced using 1/5th of the lowest positive value.

### Data availability

Datasets can be found in Supplementary Tables 1-3.

---

**Extended Data Figure 1.**
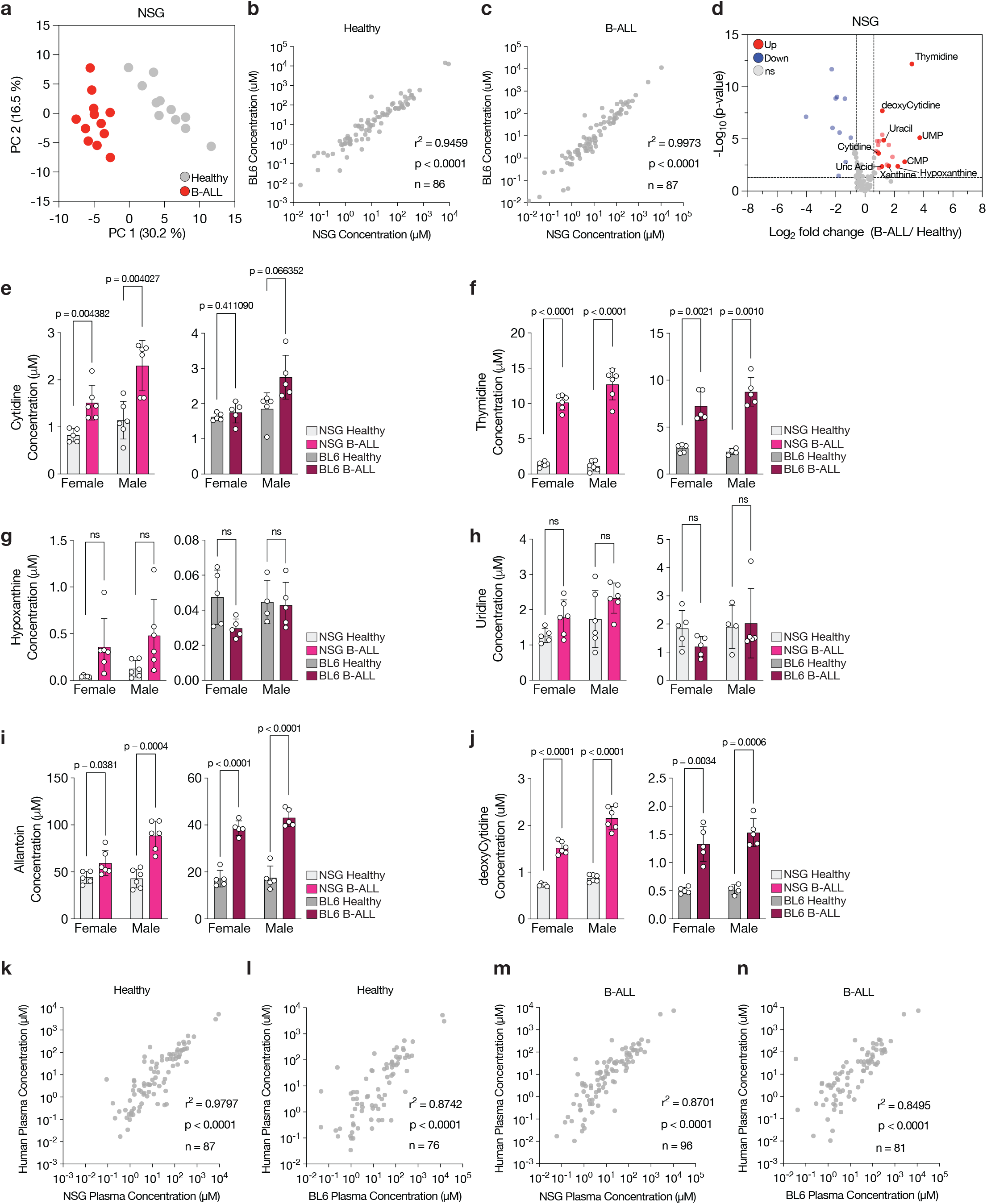
Characterization of metabolites in plasma from mice and humans with B-ALL. **a**, Principal component analysis (PCA) of metabolites quantified by LC–MS in plasma from healthy and B-ALL–bearing NSG mice. Data represent n = 11 healthy and 12 B-ALL biologically independent samples. **b–c**, Scatter plots comparing plasma metabolite concentrations measured in healthy C57BL/6J (BL6) and healthy NSG mice (b) and in BL6 and NSG mice with B-ALL (c). Each point represents the mean concentration measured for one metabolite in plasma. Data represent biologically independent samples from n = 11 (healthy NSG mice), n=10 (healthy BL6 mice; B-ALL-bearing BL6 mice), n=12 (B-ALL-bearing NSG mice). The number of metabolites measured is indicated in each panel. Pearson correlation coefficients (r) and P values from two-sided tests are shown. **d**, Volcano plot of polar metabolite concentrations in plasma from B-ALL–bearing versus healthy NSG mice. Metabolites related to nucleotide metabolism that differ between groups are labeled. Significance was defined as |log2 fold change| > 0.6 and raw P < 0.05. Data represent n = 12 diseased mice and n = 11 healthy mice. **e–j**, Plasma concentrations of individual nucleotide-related metabolites in male and female NSG and BL6 mice without or with B-ALL as indicated. Data are mean ± SD and represent n = 6 (female NSG diseased; male NSG healthy; male NSG diseased), n = 5 (female NSG healthy; female BL6 healthy; female BL6 diseased; male BL6 diseased), and n = 4 (male BL6 healthy) biologically independent mice. Significance was determined by multiple unpaired two-tailed t-tests with Welch’s correction, and P values were adjusted for multiple comparisons using the Holm–Šídák method; ns, not significant. **k–n**, Scatter plots comparing plasma metabolite concentrations between healthy pediatric patient plasma and plasma from healthy NSG (k) or BL6 mice (l), and between pediatric Ph+ B-ALL patient plasma and plasma from NSG mice (m), or BL6 mice (n) with B-ALL. Each point represents the mean concentration of one metabolite in plasma. The number of metabolites measured is indicated in each panel. Pearson correlation coefficients (r) and P values from two-sided tests are shown. Data represent biologically independent samples from n = 11 (healthy NSG mice; healthy pediatric samples), n = 10 (healthy BL6 mice; leukemic BL6 mice), n = 12 (leukemic NSG mice), and n = 5 (pediatric Ph^+^ B-ALL samples).

**Extended Data Figure 2.**
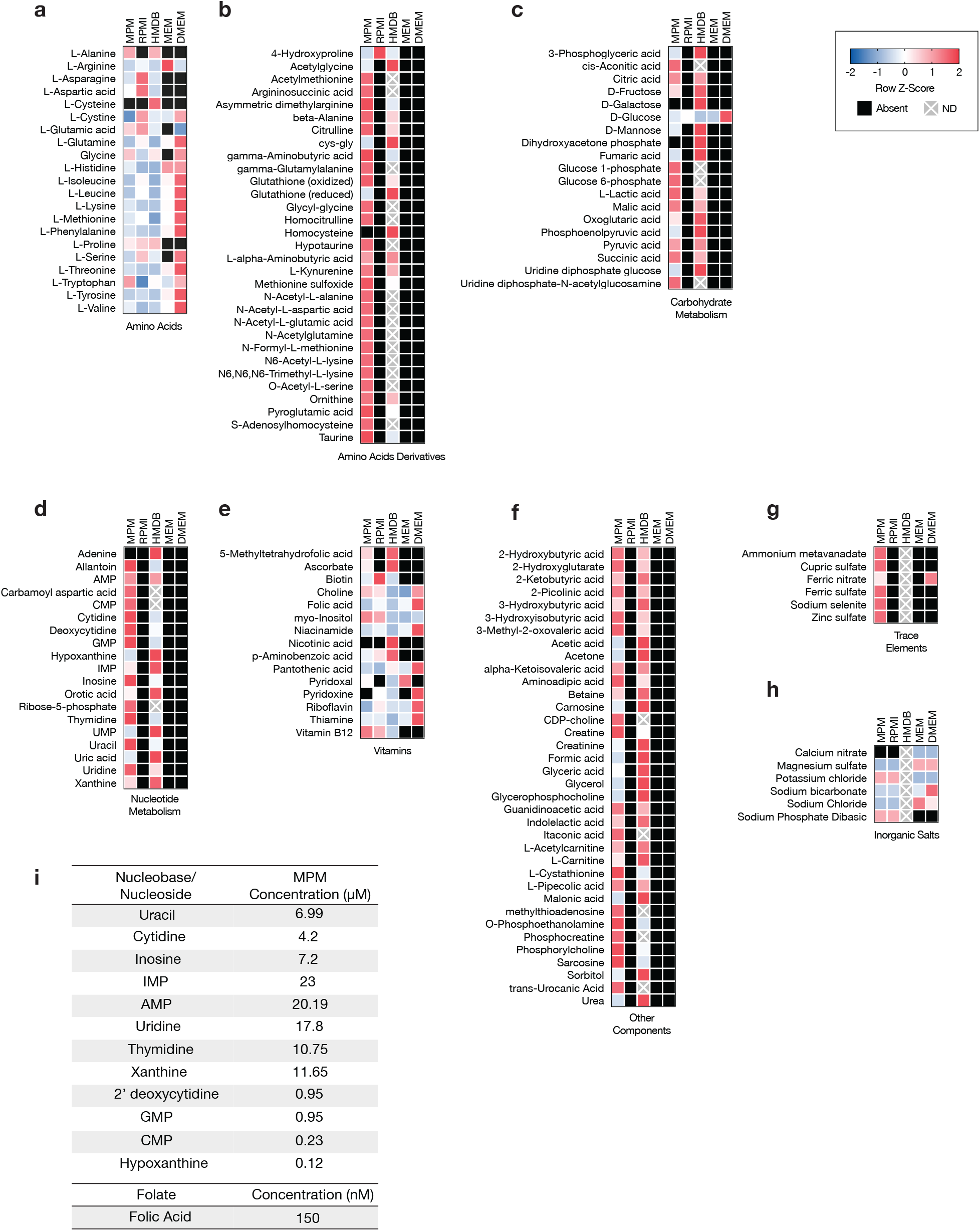
Composition of MPM compared to standard media and metabolites reported for human plasma. **a–h**, Heatmaps comparing metabolite concentrations in mouse plasma-like medium (MPM), reported for human plasma in the Human Metabolome Database (HMDB), and that used to formulated various standard culture media (RPMI, MEM and DMEM). Metabolites are grouped by metabolic pathway; values within each row were z-score normalized. Black boxes indicate metabolites absent from a medium formulation, and grey boxes with an “X” indicate metabolites with no reported concentration in HMDB (“ND”, not determined). **i**, Absolute concentrations of nucleobases, nucleosides, nucleotides and folic acid in MPM.

**Extended Data Figure 3.**
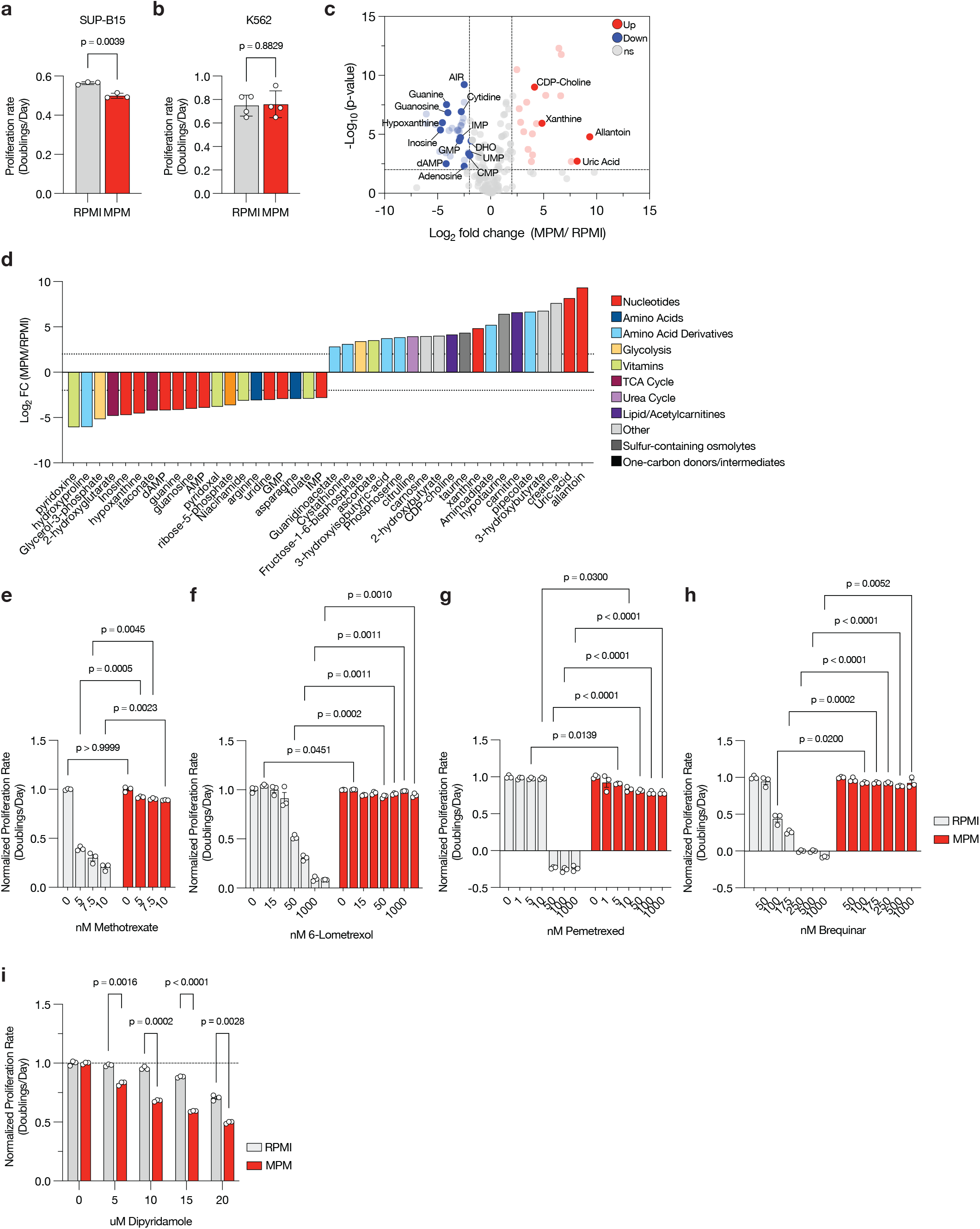
Culturing cells in MPM impacts nucleotide metabolism. **a-b**, Proliferation rates (doublings per day) of human SUP-B15 B-ALL cells (a) and K562 cells (b) cultured in RPMI or MPM. Data are mean ± SD and represent n = 3 (SUP-B15) or n = 4 (K562) biologically independent samples. Significance was determined by unpaired two-tailed t-test with Welch’s correction. **c-d**, Volcano plot (c) and pathway-level categorization (d) of log2 fold changes in intracellular metabolites measured in human SUP-B15 B-ALL cells cultured in MPM relative to RPMI for 8 days. In (c), significance was defined as |log2 fold change| > 2 and raw P < 0.01. Metabolites related to nucleotide metabolism that differ between groups are labeled. For (d), the top 20 upregulated and top 20 downregulated metabolites meeting this threshold were selected and annotated by pathway. Horizontal lines in (d) indicate |log2 fold change| = 2. Data represent n = 6 biologically independent samples. **e–h**, Proliferation rates (normalized to DMSO control) of B-ALL cells cultured in RPMI or MPM without or with the nucleotide synthesis inhibitors methotrexate (e), lometrexol (f), pemetrexed (g) or brequinar (h) as indicated. Data are mean ± SD and represent n = 3 biologically independent samples. Significance was determined by unpaired two-tailed t-tests with Welch’s correction and adjusted for multiple comparisons using the Holm–Šídák method. **i**, Proliferation rates (normalized to DMSO control) of cells cultured in RPMI or MPM without or with the nucleoside uptake inhibitor dipyridamole as indicated. Data are mean ± SD and represent n = 3 biologically independent samples. Significance was determined by unpaired two-tailed t-tests with Welch’s correction and adjusted for multiple comparisons using the Holm–Šídák method.

**Extended Data Figure 4.**
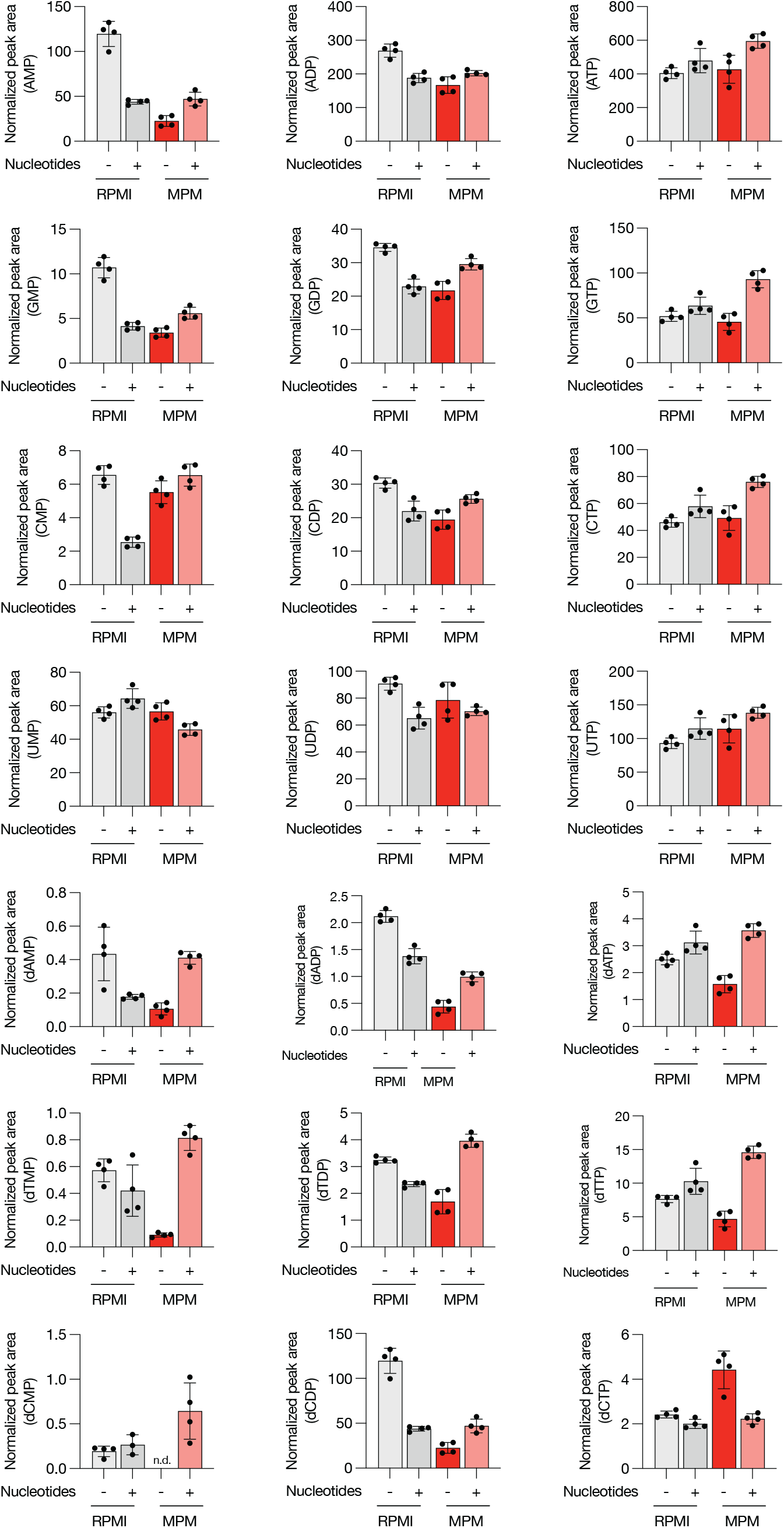
Nucleotide levels in media influence nucleotide levels in cells when cultured in either RPMI or MPM. Normalized peak areas of the indicated ribo- and deoxyribonucleotide levels measured in B-ALL cells cultured in RPMI or MPM with or without MPM-level nucleotides for 4 days. Data are mean ± SD and represent n = 4 biologically independent samples. (“n.d.”, not detected by LC-MS).

**Extended Data Figure 5.**
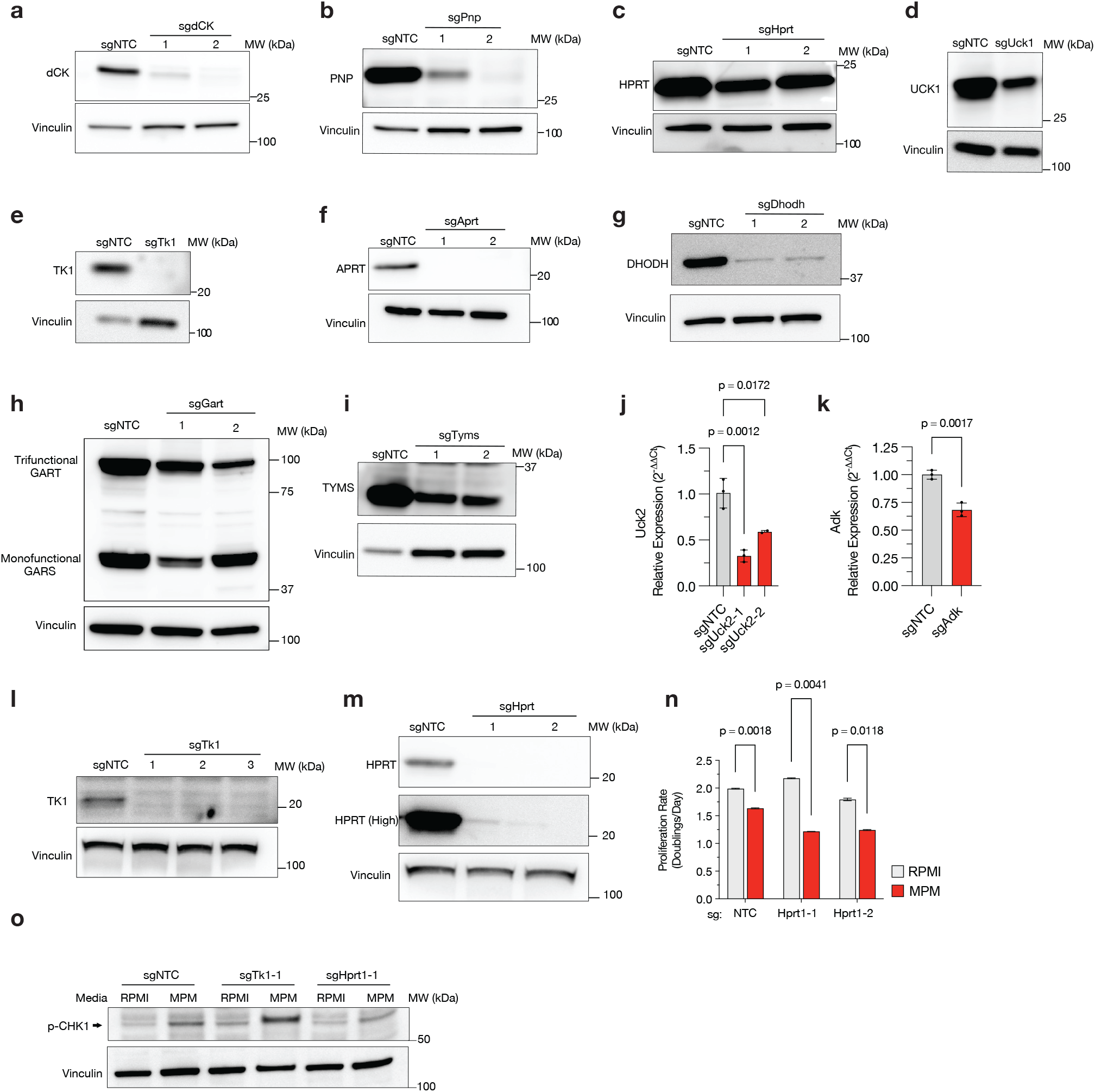
Validation of nucleotide synthesis and salvage gene knockout in B-ALL cells. **a–i**, Western blot analysis for expression of the indicated *de novo* synthesis and salvage pathway enzymes in polyclonal sgRNA-targeted B-ALL cells. Vinculin is shown as a loading control. **j-k**, Relative mRNA expression of Uck2 (j) and Adk (k) measured by RT–qPCR in sgRNA-targeted B-ALL cells compared to sgNTC controls. Data are mean ± SD and represent n = 3 biologically independent samples. **l-m**, Western blot analysis for expression of the indicated proteins in clonal sgTk1 (l) and sgHprt (m) B-ALL cell lines. Vinculin is shown as a loading control. **n**, Proliferation rates (doublings per day) of sgNTC and sgHPRT clonal B-ALL cells cultured in RPMI or MPM for 4 days. Data are mean ± SD and represent n = 2 biologically independent samples. **o**, Western blot analysis of sgNTC, sgTk1 and sgHprt clonal B-ALL cell lines cultured in RPMI or MPM for 4 days for phospho-Chk1 (P-Chk1 S345). Vinculin is shown as a loading control.

**Extended Data Figure 6.**
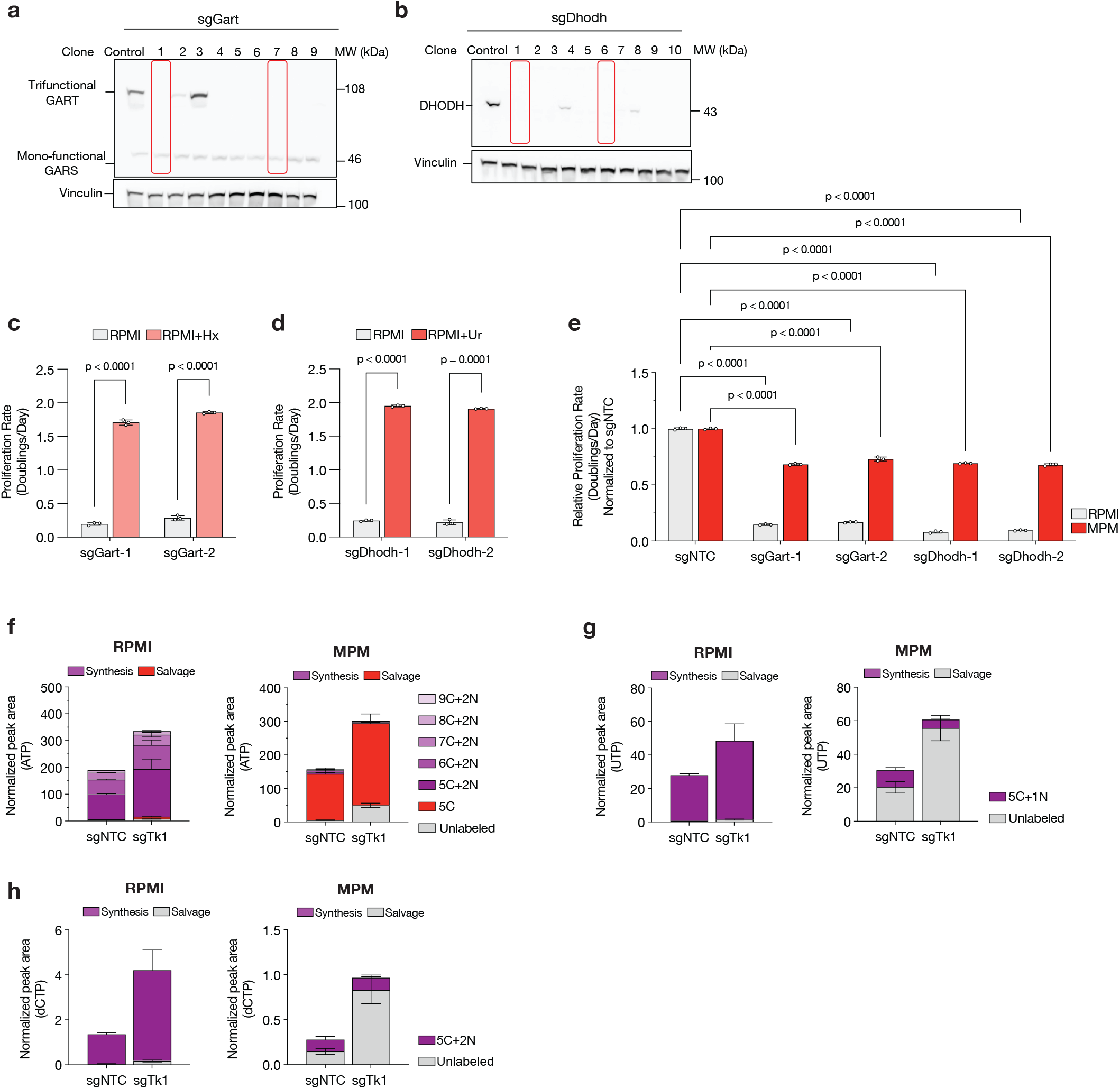
Effects of disrupting nucleotide synthesis and salvage enzymes on nucleotide metabolism and cell proliferation. **a-b**, Western blot analysis for expression of the indicated proteins in sgGart (a) and sgDhodh (b) clonal B-ALL cell lines. Vinculin is shown as a loading control. **c-d**, Proliferation rates (doublings per day) of sgGart (c) and sgDhodh(d) B-ALL clones cultured in RPMI in the presence or absence of 100 μM hypoxanthine (Hx) (c) or 100 µM uridine (Ur) (d). Data are mean ± SD and represent n = 3 biologically independent samples. Statistical significance was determined using two-tailed unpaired Welch’s t-tests, with Holm–Šidák correction for multiple comparisons. **e**, Relative proliferation rates (doublings per day, normalized to sgNTC controls) of the indicated sgGart or sgDhodh clones cultured in RPMI or MPM. Data are mean ± SD and represent n = 3 biologically independent samples. Significance was determined by ordinary one-way ANOVA followed by Holm–Šídák multiple-comparisons test. **f–h**, Isotopologue distributions of ATP (f), UTP (g) and dCTP (h) in B-ALL cells cultured for 24 h with [U-^13^C_6_]glucose and [amide-^15^N]glutamine in RPMI or MPM as indicated. Data are mean ± SD and represent n = 3 biologically independent samples.

**Extended Data Figure 7.**
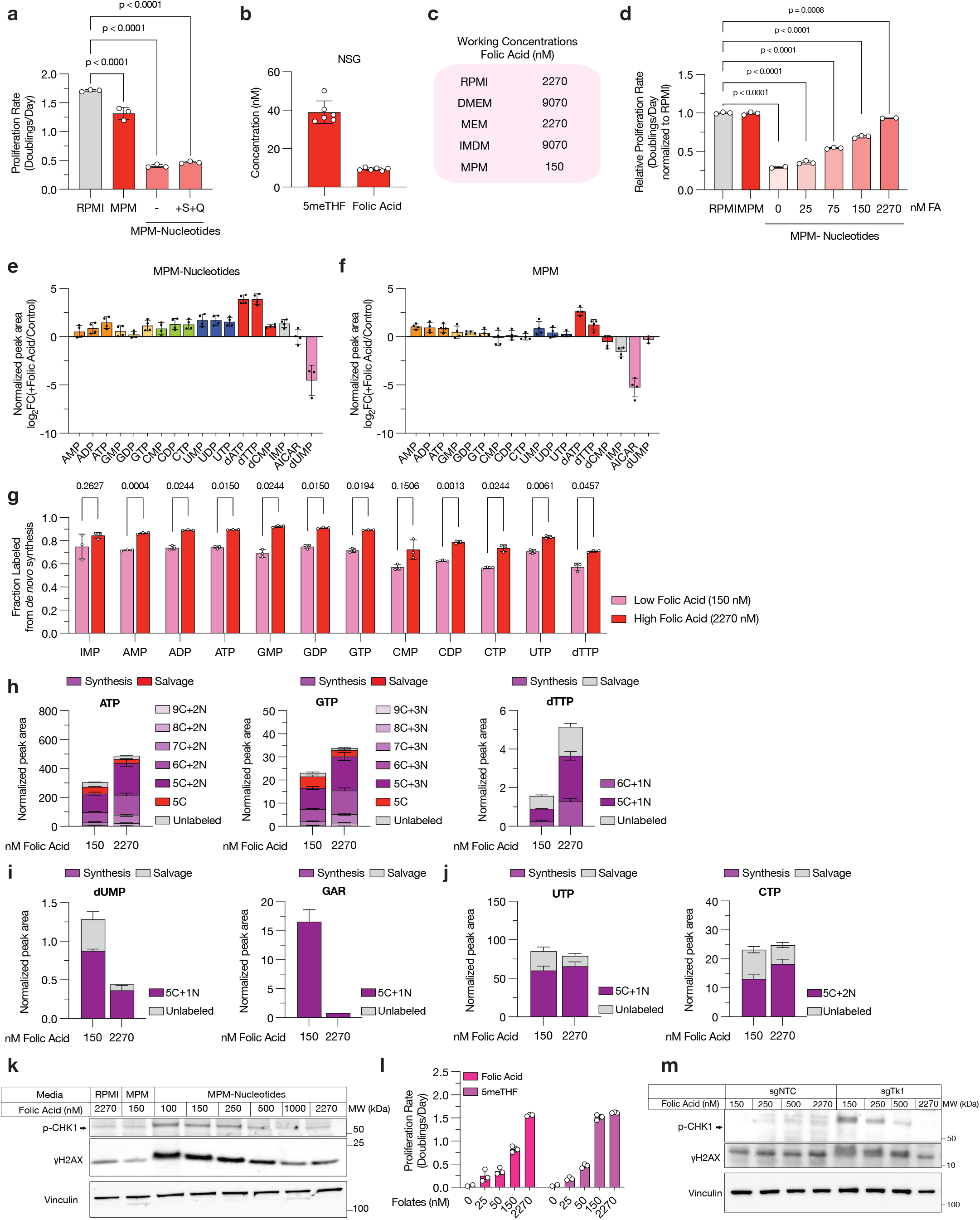
Physiological folate levels modulate *de novo* thymidylate and purine synthesis. **a**, Proliferation rates (doublings per day) of B-ALL cells cultured for 4 days in RPMI, MPM, or MPM without nucleotides supplemented without or with 286 μM serine (S) and 2mM glutamine (Q) as indicated. Data are mean ± SD and represent n = 3 biologically independent samples. Significance was determined by ordinary one-way ANOVA followed by Holm–Šídák multiple-comparisons test. *Data for RPMI, MPM, and MPM-Nuc are from the same experiment shown in Fig. 3b*. **b**, Plasma concentrations of 5-methyltetrahydrofolic acid (5meTHF) and folic acid measured in NSG mice. Data are mean ± SD and represent n = 6 biologically independent samples. **c**, Table summarizing folic acid concentrations in commonly used culture media formulations and in MPM. **d**, Relative proliferation rates (doublings per day, normalized to culture in RPMI) of SUP-B15 cells cultured for 4 days in RPMI, MPM, or MPM without nucleotides without or with the indicated concentrations of folic acid (FA). Data are mean ± SD and represent n = 3 biologically independent samples. Significance was determined by ordinary one-way ANOVA followed by Holm–Šídák multiple-comparisons test. **e-f**, Log2 fold change in levels of the indicated nucleotide species measured in B-ALL cells cultured for 4 days with 2.27 μM folic acid compared to 150 nM folic acid in MPM without nucleotides (e) or in MPM (f). Data are mean ± SD and represent n = 4 biologically independent samples. **g**, Fractional contribution of *de novo* synthesis to the indicated nucleotide pools in B-ALL cells cultured in MPM-nucleotides with 150 nM or 2.27 μM folic acid for 24 hours. *De novo* synthesis was defined as fraction with label incorporation into both ribose and base moieties. Data are mean ± SD and represent n = 3 biologically independent samples. Significance was determined by multiple unpaired two-tailed t-tests with Welch’s correction and adjusted for multiple comparisons using the Holm–Šídák method. **h–j**, Isotopologue distributions and normalized peak areas of indicated metabolites in B-ALL cells cultured with [U-^13^C_6_]glucose and [amide-^15^N]glutamine in MPM-Nucleotides media with 150 nM or 2.27 μM folic acid for 24 hours. Data are mean ± SD and represent n = 3 biologically independent samples. **k**, Western blot analysis of phospho-Chk1 (P-Chk1 S345) and *γ*H2AX levels in B-ALL cells cultured in RPMI, MPM, or MPM without nucleotides supplemented with the indicated folic acid concentrations. Vinculin is shown as a loading control. **l**, Proliferation rates (doublings per day) of B-ALL cells cultured in RPMI formulated with the indicated concentrations of folic acid or 5-methyltetrahydrofolic acid (5meTHF) as indicated. Data are mean ± SD and represent n = 3 biologically independent samples. **m**, Western blot analysis of phospho-Chk1 (P-Chk1 S345) and *γ*H2AX in sgNTC and sgTk1 clonal B-ALL cells cultured in MPM with the indicated folic acid concentrations. Vinculin is shown as a loading control.

**Extended Data Figure 8.**
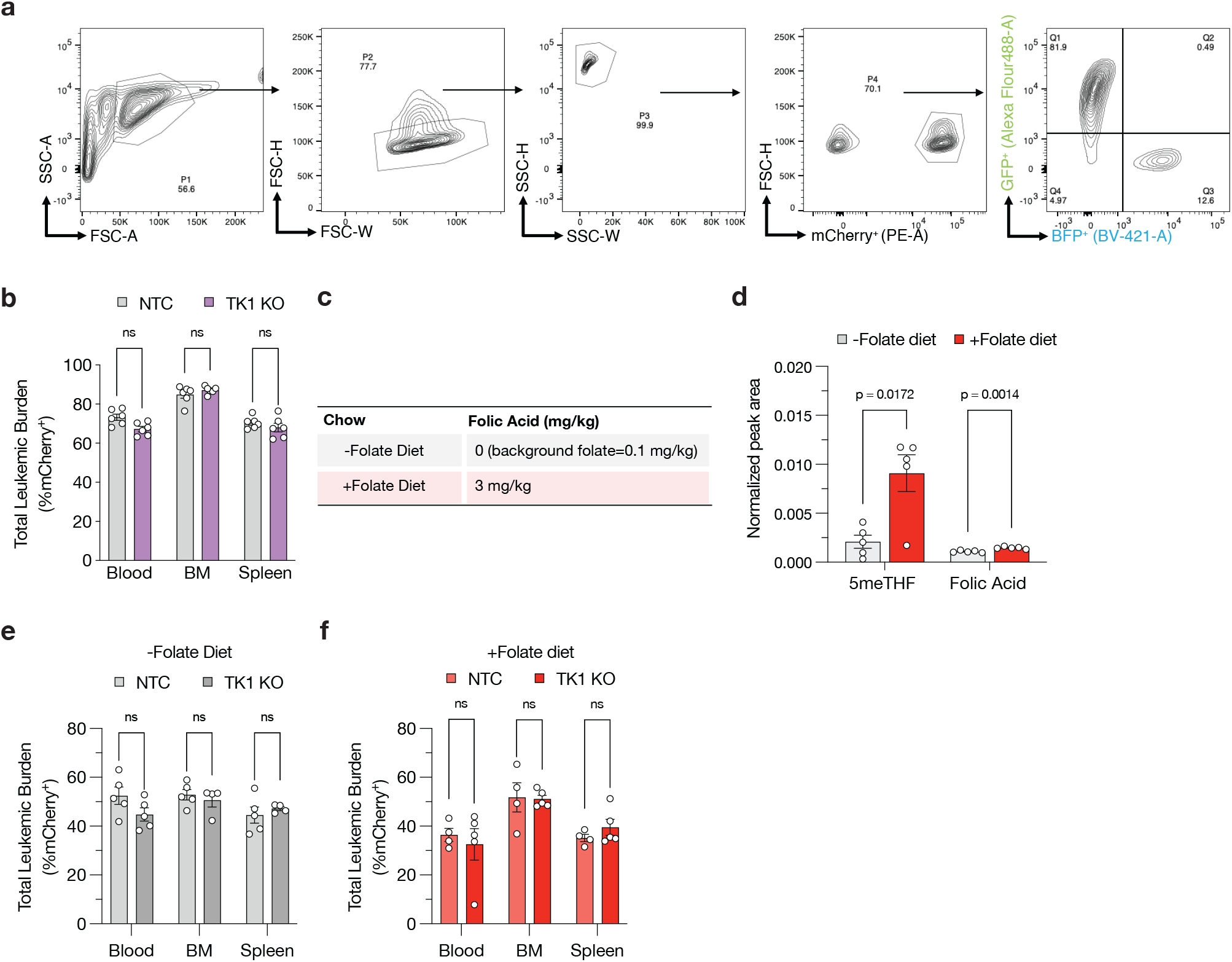
Dietary folate manipulation modulates TK1 knockout B-ALL proliferation *in vivo*. **a**, Flow cytometry gating strategy for *in vivo* competition experiments described in Fig. 4. **b**, Percent mCherry^+^ B-ALL cells measured from the indicated tissues in the experiment shown in Fig. 4a-b. Data are mean ± SEM and represent n = 6 biologically independent male NSG mice. Significance was determined by multiple unpaired two-tailed t-tests with Welch’s correction and Benjamini–Krieger–Yekutieli FDR correction. **c**, Table summarizing composition of folate-deplete (−Folate) or folate-replete (+Folate) diets. **d**, Plasma folate levels measured in mice maintained on folate-free (-Folate) or +folate (3 mg kg^−1^) diets for 4 weeks. Data are mean ± SEM and represent n = 5 biologically independent male NSG mice. Statistical significance between groups was determined using two-tailed unpaired Welch’s t-tests with Holm–Šídák correction for multiple comparisons. 5meTHF=5-methyltetrahydrofolic acid. **e-f**, Percent mCherry^+^ B-ALL cells measured from the indicated tissues in the experiment shown in Fig. 4c–f. (e), −Folate diet; (f), +Folate diet. Data are mean ± SEM and represent n = 5 (NTC −Folate; TK1 knockout (KO) −Folate; TK1 KO +Folate) and n = 4 (NTC +Folate) biologically independent female NSG mice. Significance was determined by multiple unpaired two-tailed t-tests with Welch’s correction and Benjamini–Krieger–Yekutieli FDR correction.

**Extended Data Figure 9.**
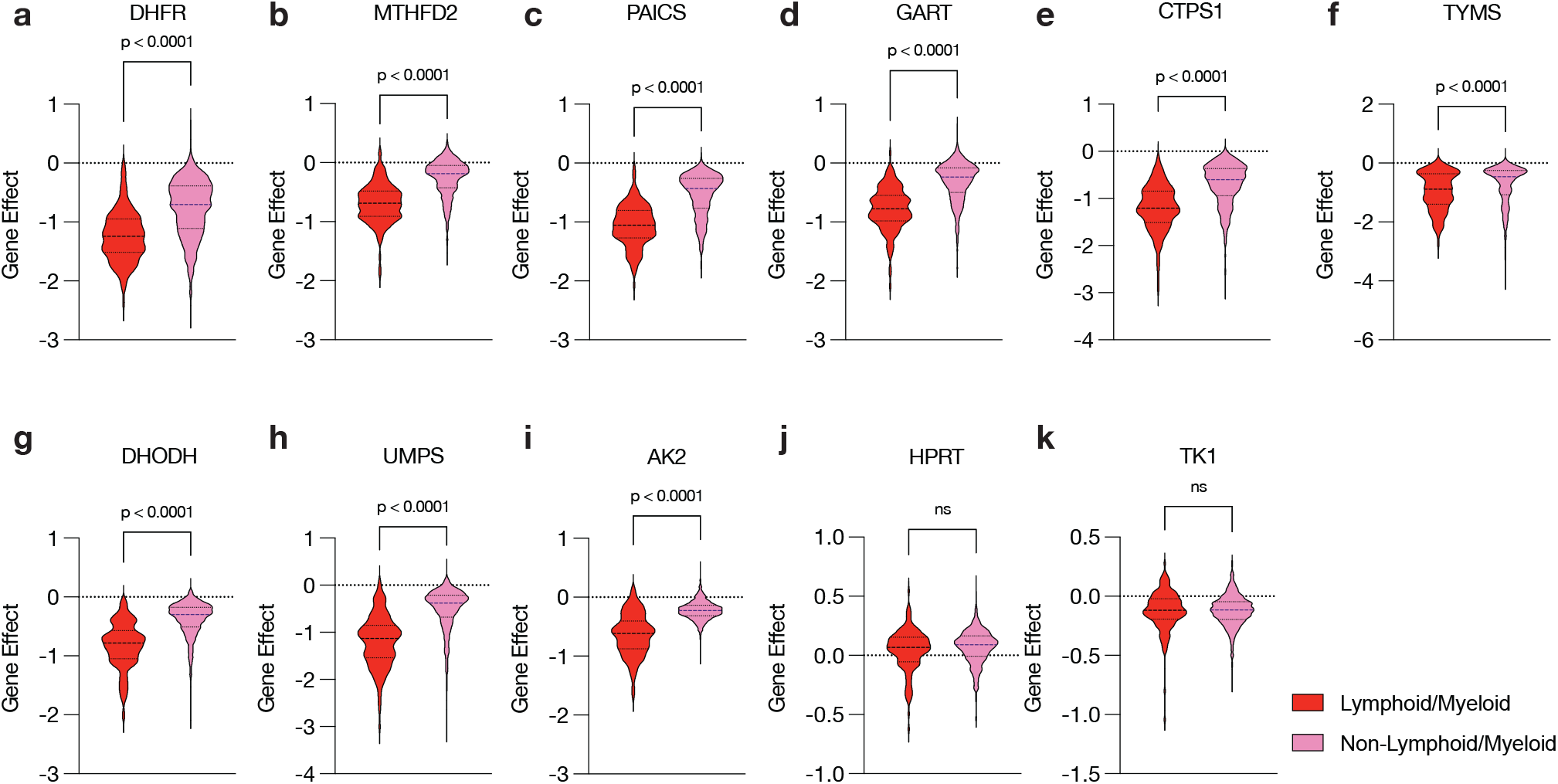
Culture in standard media may miss dependency of hemopoietic cancer cells on nucleotide metabolism genes. DepMap CRISPR gene dependency (Chronos, release 25Q3+) scores for *de novo* nucleotide synthesis and nucleotide salvage pathway genes across hematopoietic (lymphoid and myeloid) versus non-hematopoietic cell lines. Significance was determined by unpaired two-sided Mann–Whitney U test; ns: not significant.

